# Helicase-Like Functions in Phosphate Loop Containing Beta-Alpha Polypeptides

**DOI:** 10.1101/2020.07.30.228619

**Authors:** Pratik Vyas, Olena Trofimyuk, Liam M. Longo, Fanindra Kumar Deshmukh, Michal Sharon, Dan S. Tawfik

**Affiliations:** Department of Biomolecular Sciences, Weizmann Institute of Science, Rehovot, Israel; Tokyo Institute of Technology, Earth-Life Science Institute, Tokyo, Japan and Blue Marble Space Institute of Science

## Abstract

The P-loop Walker A motif underlies hundreds of essential enzyme families that bind nucleotide triphosphates (NTPs) and mediate phosphoryl transfer (P-loop NTPases), including the earliest DNA/RNA helicases, translocases and recombinases. What were the primordial precursors of these enzymes? Could these large and complex proteins emerge from simple polypeptides? Previously, we showed that P-loops embedded in simple βα repeat proteins bind NTPs, but also, unexpectedly so, ssDNA and RNA. Here, we extend beyond the purely biophysical function of ligand binding to demonstrate rudimentary helicase-like activities. We further constructed simple 40-residue polypeptides comprising just one β-(P-loop)-α element. Despite their simplicity, these P-loop prototypes confer functions such as strand separation and exchange. Foremost, these polypeptides unwind dsDNA, and upon addition of NTPs, or inorganic polyphosphates, release the bound ssDNA strands to allow reformation of dsDNA. Binding kinetics and low-resolution structural analyses indicate that activity is mediated by oligomeric forms spanning from dimers to high-order assemblies. The latter are reminiscent of extant P-loop recombinases such as RecA. Overall, these P-loop prototypes comprise a plausible description of the sequence, structure and function of the earliest P-loop NTPases. They also indicate that multifunctionality and dynamic assembly were key in endowing short polypeptides with elaborate, evolutionarily relevant functions.

**Significance statement:** It is widely assumed that today’s large and complex proteins emerged from much shorter and simpler polypeptides. Yet the nature of these early precursors remains enigmatic. We describe polypeptides that contain one of the earliest protein motifs, a phosphate-binding loop, or P-loop, embedded in a single beta-alpha element. These P-loop prototypes show intriguing characteristics of a primordial world comprised of nucleic acids and peptides. They are ‘generalists’ capable of binding different phospho-ligands, including inorganic polyphosphates and single-stranded DNA. Nonetheless, in promoting double-stranded DNA unwinding and strand-exchange they resemble modern P-loop helicases and recombinases. Our study describes a missing link in the evolution of complex proteins – simple polypeptides that tangibly relate to contemporary P-loop enzymes in sequence, structure and function.

## Introduction

Protein machines, such as ATP synthetases, helicases or RecA recombinases, perform remarkable actions and are immensely complex^1,2^. Among other factors, complexity is manifested in the concerted action of multiple functional elements, the productive orchestration of which demands large, well-defined structures. This complexity is, however, at odds with the assumption that proteins emerged by duplication and fusion of relatively short and simple polypeptides^3,4^. Albeit, these polypeptides predated the last universal common ancestor (LUCA) and as such cannot be reconstructed by conventional phylogenetic methods nonetheless, despite nearly 4 BY of evolution, and extensive diversification of sequence, structure and function, these ancestral polypeptides are traceable by virtue of being the most conserved and functionally essential motifs of modern-day proteins^5^. Perhaps the most widely spread motif is the Walker-A P-loop^6^. Defined as GxxxxGK(T/S) or GxxGxGK, this motif underlies the most abundant and diverse protein class: the P-loop NTPases^6–9^. Structurally, the P-loop NTPase domain comprises a tandem repeat of at least five β-(loop)-α elements arranged in a αβα 3-layer sandwich architecture^7,8,10^. The loops connecting the C-termini of the β-strands to the N-termini of the following α-helices comprise the active site, while short loops link the β-(loop)-α elements to one another (here, unless stated otherwise, loops refer to the former). The Walker A P-loop is the key element mediating NTP binding and catalysis, and it uniformly resides within the first β-(loop)-α element, henceforth designated as βl-(P-loop)-α1.

P-loop NTPases are among the most ancient enzyme families – if not the most ancient enzyme family itself – and the first P-loop NTPase likely emerged at the transition from the RNA world to the primordial RNA-protein world^5,11–13^. Binding of phospho-ligands, foremost ATP and GTP, is the founding function of P-loop NTPases^14^. In agreement with early emergence, the key to phospho-ligand binding is not only the P-loop, but also a ‘crown’ of hydrogen bonds realized by the end of the Walker A motif (GK(T/S) that comprises the first turn of α1 (Refs.^14,15^). It has been accordingly hypothesized that polypeptides comprising the P-loop and its flanking secondary structural elements, namely, a β-(P-loop)-α segment, were the seed from which modern P-loop NTPases emerged^5,16–18^. However, that relatively short polypeptides can confer an evolutionarily meaningful function is far from obvious. Polypeptides generally lack structural volume and complexity, and cannot, in principle, align multiple functional elements as do intact protein domains^19^. Hence, identifying functional polypeptides, and especially polypeptides that are assigned as the seeding elements of the earliest enzyme families, is of paramount importance to our understanding of protein evolution.

Previously, we showed that an ancestral β1-(P-loop)-α1 motif, inferred via phylogenetic analysis of all known P-loop NTPase families, could be grafted onto a simple repeat scaffold comprised of four consecutive βα elements^16^. This grafting resulted in simple proteins that contained two P-loops. These bound via the P-loop motif ATP and GTP, but rather unexpectedly, also RNA and ssDNA. It is therefore likely that the primordial P-loop was a multifunctional phospho-ligand binder that could function in the absence of other auxiliary residues^16^. In the contemporary P-loop NTPases, single-stranded (ss) DNA binding is mostly conferred by specialized domains^20–23^ that are distinct from the ATP binding P-loop^24^. And yet, in support of the P-loop’s multifunctionality, we searched for and revealed examples of ssDNA binding mediated by the Walker A P-loop (detailed in ‘Discussion’). Further, many of the P-loop NTPases families that date back to LUCA are involved in RNA/DNA remodeling, including helicases, RecA like recombinases, and translocases^25^. ATP synthetase, for example, is thought to have emerged from such nucleic acid-remodeling P-loop NTPases^26^.

That ssDNA binding can be mediated by our P-loop prototypes, and the possible origins of P-loop NTPases as nucleic acid remodelers, guided us to extend our previous study^16^ beyond the realm of ligand binding *per se.* We now *(i)* examine the potential of P-loop prototypes to exert, biologically and evolutionarily relevant helicase-like functions, *i.e.,* to mediate unwinding of dsDNA and strand exchanges, and (*ii*) identify the minimal structural context in which the P-loop can confer these relevant functions.

## Results

### P-loop prototypes mediate strand separation

dsDNA undergoes spontaneous local fluctuations referred to as “DNA breathing,”^27,28^ thus giving access to ssDNA binding proteins^29^. For instance, T4 helicases bind to and stabilize these transient open regions to unwind dsDNA^28^ — a mode of unwinding also referred to as “passive unwinding”^30^. To test the potential of P-loop prototypes to facilitate such passive unwinding, we used a standard fluorescence-based helicase assay dubbed a molecular beacon^31^ (**Figure 1A**). This assay uses a dsDNA segment comprised of a “beacon oligo” with a fluorophore at the 5’ end and a quencher at the 3’ end (the beacon sense strand) hybridized to an “unlabeled” complementary antisense-strand. Upon addition of a protein that promotes strand separation, owing to its complementary ends, the beacon strand collapses into an intramolecular hairpin (stem-loop structure). This brings together the fluorophore and the quencher, thus quenching the beacon’s fluorescent signal. As shown later, this stem-loop product is likely stabilized by the fluorophore-quencher interaction^32^ thus providing a simple way of monitoring DNA unwinding^31^.

**Figure 1.**
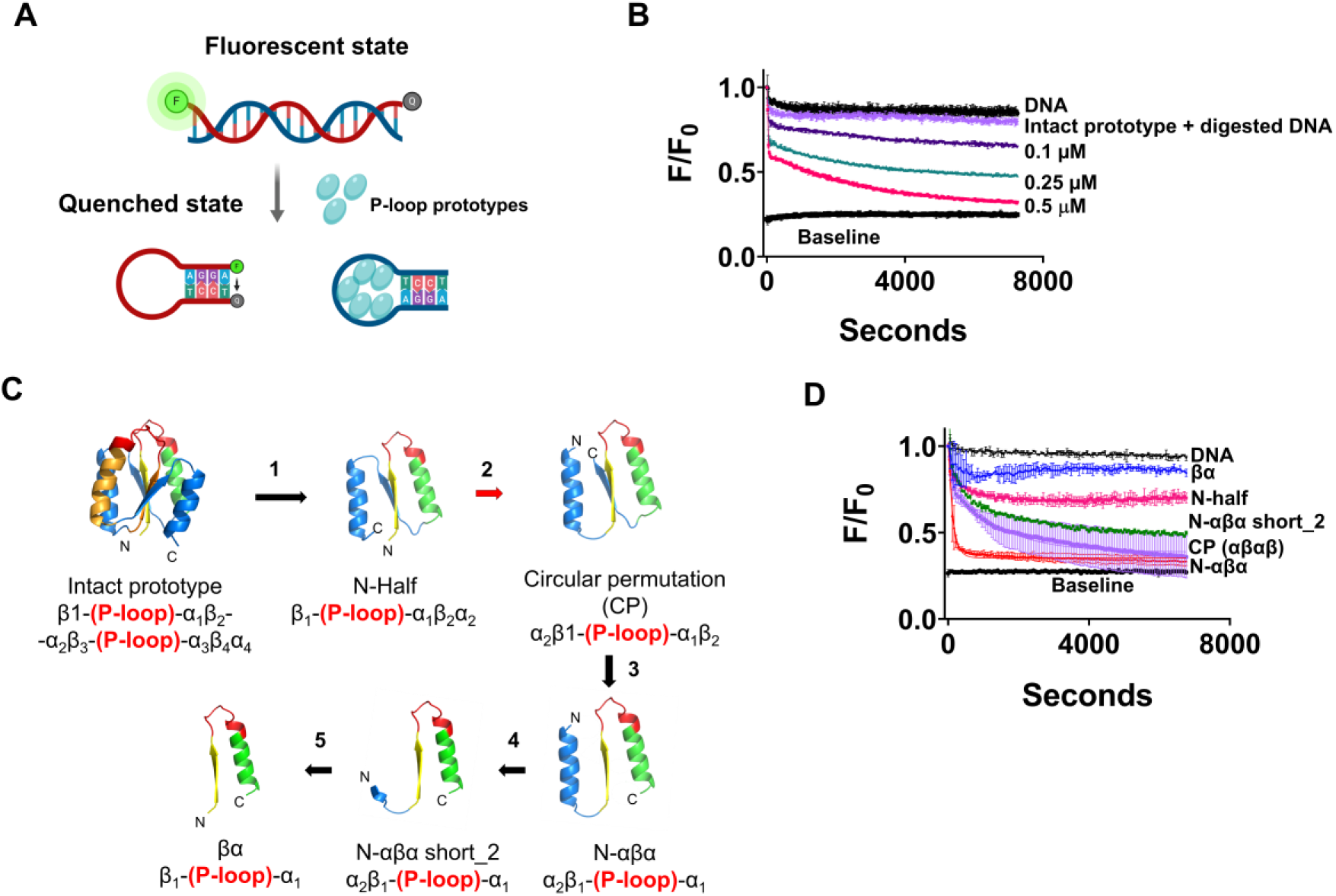
The molecular beacon assay reports the strand separation of P-loop prototypes. **A.** Simplified schematic of the strand-separation molecular beacon assay^31^. In the initial dsDNA state, the quencher of the beacon strand is held apart from the fluorophore resulting in energy transfer (high fluorescence). The preferential binding of P-loop prototypes to ssDNA induces strand separation, allowing the beacon strand to assume a hairpin state wherein the fluorophore is quenched. The oligos used in this assay, and the assays described in the subsequent figures are listed in **Supplementary Table S1**. **B.** A representative strand separation experiment using the intact 110 residue P-loop prototype (D-Ploop; Ref^16^). Strand separation is reported by the change in FRET signal (i.e. fluorescence quenching in our experimental set-up) upon addition of the P-loop prototype at increasing protein concentrations. All assays were performed with 5 nM beacon dsDNA, in 50 mM Tris (pH 8), at 24 °C. Shown are normalized F/F_0_ values, whereby the initial fluorescence of beacon dsDNA prior to protein addition takes the value of 1. Baseline represents the signal of the fully quenched hairpin beacon. Digested DNA represents the signal upon addition of 0.5 μM P-loop prototype to the beacon dsDNA pretreated with Benzonase nuclease. Traces were fitted to a biphasic exponential decay model (described in ‘**Supplementary Information**’ section) and the apparent rate constants are given in **Supplementary Table S4**. **C.** New P-loop prototypes were constructed by systematic truncation and circular permutation of the intact prototype. The numbered arrows indicate sequential steps in the engineering of new constructs as follows: *1.* Truncation of the intact 110 residue prototype into half^16^ (N-half indicates N-terminal half). *2.* Circular permutation (red arrow) of N-half to a construct with ‘αβαβ’ architecture. *3.* Truncation of the C-terminal β-strand to give N-αβα. *4-5.* Incremental truncations of N-terminal helix of N-αβα down to a βα fragment. The structural models indicate the ancestral P-loop element in yellow (β1), red (the Walker P-loop) and green (α1), while the remaining parts are in blue. **D**. Strand separation by truncated P-loop prototypes, at 1 μM protein concentration, and under the stringent condition (with 100 mM NaCl; other assay conditions as in panel A). The lines represent the average from two to six independent experiments and the bars represent the S.D. values.

For the initial round of experiments, the originally described 110 amino acid protein that presents two P-loops (D-Ploop^16^), hereafter referred to as the ‘intact prototype’, was used. This intact prototype bound ssDNA preferably over double strand (ds) DNA^16^. As shown later, the P-loop prototypes also exhibit preference for TC-rich ssDNA while GA-rich sequences are barely bound. Accordingly, the dsDNA beacon comprised a TC-rich unlabeled antisense strand to promote binding and strand separation, while the beacon sense strand was GA-rich to avoid interference with the stem-loop formation and non-specific quenching due to protein binding (oligonucleotide sequences are provided in **Supplementary Table S1**). When added to the beacon dsDNA, the intact prototype induced strand separation as indicated by the decrease in fluorescence (**Figure 1B**). The P-loop prototype did not induce quenching of the fluorophore itself as indicated by a negligible change in fluorescence intensity with digested beacon dsDNA (**Figure 1B**; nonetheless, this non-specific quenching was subtracted to give the quenching traces reported below, see Methods). Strand-separation, as reported by this beacon dsDNA, was applied for testing various fragments of the intact P-loop prototype as described in the next section. Subsequently, using the most active fragment, the kinetics and mechanism of strand-separation were further investigated as described below (see *The N-αβα P-loop prototype shows avid and cooperative strand separation).*

### Structural minimization of the P-loop prototype

The intact P-loop prototypes were based on an ‘ideal fold’^33^ that reproduces the 3-layered αβα sandwich architecture of P-loop NTPases^7,8,10,25^. They are composed of four βα elements with the ancestral β-(P-loop)-α element replacing the first and third elements^16^. We asked whether the intact prototype could be truncated while retaining its function. In other words, what is the minimal ‘stand-alone’ P-loop fragment that possesses phospho-ligand binding and strand-separation activity?

The repeat nature of our prototypes allowed multiple options for single P-loop constructs with varied structural topologies. In the first step, following our original report, the first half of the intact P-loop prototype (N-half) was tested^16^. Other ‘half’ constructs comprising two strands and two helices were also tested (**Figure 1C**, **Supplementary Figure S1**; DNA and amino acids sequences of all constructs are provided in **Supplementary Tables S2 and S3**). Secondly, we used circular permutation to generate half fragments that possessed a αβαβ architecture (**Figure 1C**). All of these constructs, and especially the circular permutants, showed high expression and purification yields (via a His-tag and Ni-NTA; **Supplementary Figure S2A**). Nonetheless, to maintain solubility at high protein concentrations, an osmolyte such as L-arginine was needed. We then truncated the circularly permuted ‘half’ constructs further to obtain the minimal constructs described below (**Figure 1C, Supplementary Figure S1**).

All ‘half’ constructs, and most of their truncated versions, mediated strand-separation to some degree (**Supplementary Figure S2B**). However, nucleic acid binding is known to be highly sensitive to ionic strength, and indeed the activity of most constructs was diminished in the presence of 100 mM NaCl (**Figure 1D, Supplementary Figure S2C**). We therefore set our assay conditions at 100 mM NaCl as a benchmark for robust activity. Intriguingly, under this stringent binding condition, a truncated variant dubbed ‘N-αβα’, comprising the ancestral β-(P-loop)-α element with one additional helix that precedes it, showed the most efficient and rapid strand separation (**Figure 1D**, red trace).

### The minimal P-loop fragment

Can the N-αβα construct be further shortened? Given its polar nature, the primary role of N-terminal helix might be to enhance protein solubility (**Figure 1C, steps 4-6, Supplementary Table S3**). Indeed, partial truncations of this helix showed very similar activity to the intact N-αβα (N-αβα short_2, **Figure 1D**, green trace), while its complete truncation resulted in loss of the strand-separation activity (βα, **Figure 1D**, blue trace). Nonetheless, this shortest construct, which in effect comprises a β-(P-loop)-α fragment, tended to co-purify with nucleic acids, suggesting that it retained some binding capability. The functional prototype N-αβα short_2 contains only a short N-terminal segment (KRRGV) linked to the β-(P-loop)-α fragment by a short linker (GSG; **Supplementary Table S3**). It therefore appears that given a charged solubility tag, a single β-(P-loop)-α fragment can confer strand-separation activity. Also, worth noting is that N-αβα possesses low sequence complexity – it is made of only 12 amino acid types. Ten of these are considered abiotic amino acids, plus two cationic amino acids, Lys and Arg (excluding the 6xHistag for purification, and a tryptophan introduced for determining concentration by absorbance at 280 nm; **Supplementary Table S3**).

Overall, we found that the core β-(P-loop)-α motif can be placed in a variety of structural contexts, with varying strand topologies, and yet retain, or even show improved biochemical function (**Figure 1D**, **Supplementary Figures S1 and S2**). Given its minimal size and sequence complexity, its high expression, purity, and foremost its high strand separation activity, the N-αβα construct was employed as the representative P-loop prototype for the studies described below.

### The N-αβα P-loop prototype shows avid and cooperative strand separation

Strand separation was dependent on the concentration of P-loop prototypes, in terms of both the rate and amplitude of fluorescence decay. At higher concentrations, unwinding was faster and complete (*i.e.*, reached baseline) whereas at lower concentrations unwinding was slower and partial (**Figure 2A, Supplementary Tables S4 and S5**). Isotherms obtained by plotting the endpoint fluorescence-decay values *versus* protein concentration indicated that strand separation is highly cooperative (sigmoidal curves and high Hill’s coefficient) and occurs with apparent K_D_ values in the sub μM range (**Figure 2C, Supplementary Table S6**). The unwinding kinetics are complex since the dependence of the apparent rate constants on protein concentration was not linear (**Figure 2B**). Thus, strand separation appears to be a multiple step process. Specifically, the rate determining step changes non-linearly with protein concentration and at > 0.5 μM becomes essentially concentration independent (**Figure 2B**). As shown later, both of these phenomena (cooperative isotherms and complex kinetics) are in agreement with changes in quaternary structure upon ligand binding. Mutations in the key P-loop residues of N-αβα reduced both the rate and the amplitude of fluorescence decay, and accordingly showed loss of binding to ssDNA by ELISA (**Supplementary Figure S3**).

**Figure 2.**
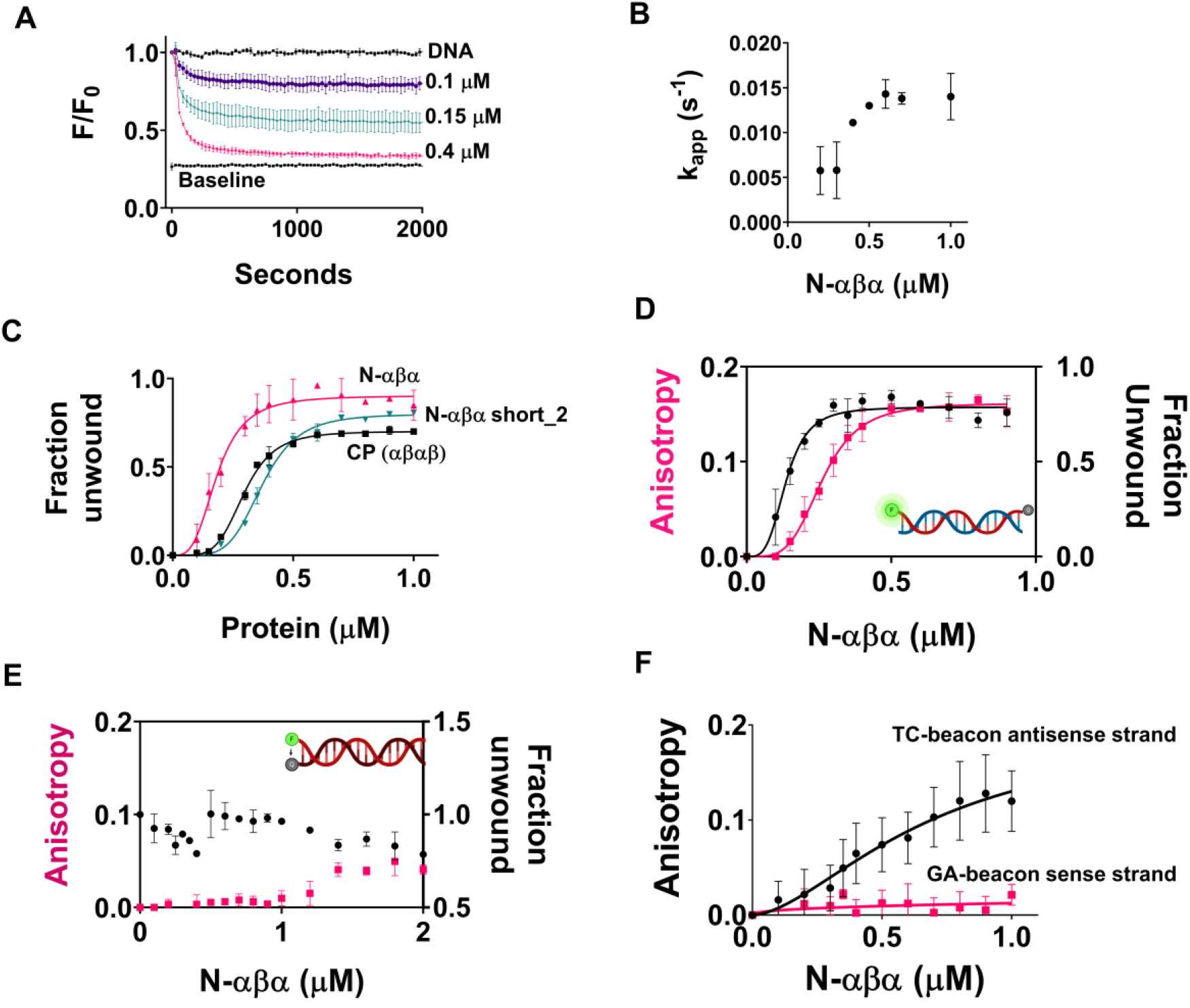
Binding affinities and kinetics of strand-separation by P-loop prototypes. **A.** A representative strand-separation experiment with varying concentrations of N-αβα (the beacon dsDNA is described in **Figure 1A**, and assay conditions are as in **Figure 1D**). The values shown are the average from 2 to 6 independent experiments and the bars represent the S.D. values (the number of experiments in the subsequent panels is denoted as *n*). **B.** Traces shown in panel *A* were fitted to a one phase exponential decay model (**Eq. 1**, **Supplementary Information**) and the apparent rate constants were plotted against protein concentration (**Supplementary Table S5)**. **C.** Binding isotherms of P-loop prototypes (for their topology, see **Figure 1C**). End-point F/F_0_ values (two hours), from strand separation experiments (*e.g*. panel A) were normalized (fluorescence of the starting beacon dsDNA equals 0, and of the fully quenched ssDNA hairpin equals 1) to derive the relative fraction of unwound dsDNA, and were then plotted vs. protein concentration (n = 2 to 6; error bars represent SD values). **D.** Simultaneous monitoring of the changes in fluorescence intensity and in anisotropy of the beacon dsDNA. Fluorescence polarization and quenching were monitored for two hours after incubating the beacon dsDNA with varying concentrations of N-αβα. Changes in anisotropy (pink trace) and in the fraction unwound (black trace; as in panel *C*) were measured as described in *Methods* (n = 4 to 8; error bars represent SD values). Shown here are two-hour end-point values plotted against N-αβα concentration. **E**. As above but with a dsDNA construct having the fluorophore and quencher on the opposite strands (here, strand separation should result in an increase in fluorescence rather than decrease as in panel *D* and in all other beacon assays; n = 4 to 8; error bars represent SD values). **F.** Binding to the individual strands of the dsDNA beacon as monitored by fluorescence anisotropy assay. Fluorescence polarization was monitored for two hours after incubation of the GA-beacon sense strand and the TC-beacon antisense strand, with varying concentrations of N-αβα. Changes in anisotropy were measured as described in *Methods* (n = 4 to 8; error bars represent SD values). Shown here are two-hour end-point values for GA-beacon sense strand (pink trace) and the TC-beacon antisense strand (black trace) plotted against N-αβα concentration.

Unwinding of the beacon dsDNA is likely driven by the binding preference of the P-loop prototypes for ssDNA over dsDNA. This preference was originally observed with the intact prototype using an ELISA-like assay (with immobilized DNA and detection of binding with antibodies to the 6xHis tag)^16^. This result was confirmed here, also with the N-αβα fragment and the same oligonucleotides used in the beacon assay (**Supplementary Figure S4**). However, the ELISA assay exhibited high background. Thus, to more accurately measure binding to ss-versus ds-DNA, and to gain further insight into the mechanism of strand separation, we employed an assay, following a previously described setup^34^, to simultaneously measure strand separation by fluorescence quenching (as in the beacon assay, **Figure 1**) and binding to the fluorescently labelled DNA strand(s) by fluorescence anisotropy.

We first tested the beacon dsDNA. Upon addition of the N-αβα prototype, a concentration dependent change in anisotropy was observed that was concomitant with the quenching signal (**Figure 2F**). The change in anisotropy occurs in parallel with quenching suggesting that strand separation and binding occur in a concerted manner, and that the anisotropy signal reports binding to ssDNA; or in other words: the dsDNA bound states do not accumulate. Indeed, under the conditions applied here the P-loop prototypes do not exhibit dsDNA binding, as confirmed with a dsDNA construct in which the quencher resides on the complementary strand. This construct should show an increase in fluorescence upon strand separation (as opposed to the decrease observed with the beacon dsDNA). However, no such increase was observed upon prototype addition (**Figure 2E**) likely due to the to the fluorophore-quencher interaction inducing higher duplex stability^32^ (also indicated by strand exchange assays described below). Accordingly, this construct also showed a negligible change in anisotropy (**Figure 2E**).

The second principle governing dsDNA unwinding is the selectivity in binding the TC-rich antisense strand relative to the GA-rich sense strand of the beacon dsDNA. This selectivity was observed by ELISA (**Supplementary Figure S4**) as well as by anisotropy (**Figure 2F**, **Supplementary Table S7**).

Finally, N-αβα does not quench the fluorescently labeled dsDNA in the absence of the quencher containing complementary strand (**Supplementary Figure S5**). This rules out the posibility that fluorescence quenching is driven by spurious interactions of the P-loop prototype with the fluorophore.

### The P-loop prototypes facilitate strand exchange

Next, we assessed the potential of N-αβα to mediate strand-exchange – namely to accelerate the rate of exchange between a duplex bound ssDNA and an identical competing strand added in excess. We started with the dsDNA construct which, due to higher stability, did not show strand-separation (**Figure 2E; Figure 3A**). This construct also showed no exchange when mixed with 100-fold excess of the unlabeled complementary strand, not even after 24 hr. incubation (**Figure 3B**, pink bar). However, addition of N-αβα led to an increase of fluorescence indicating the displacement of the quencher strand by the unlabeled one, in a concentration and time dependent manner (6-24 hr.; **Figure 3B**).

**Figure 3.**
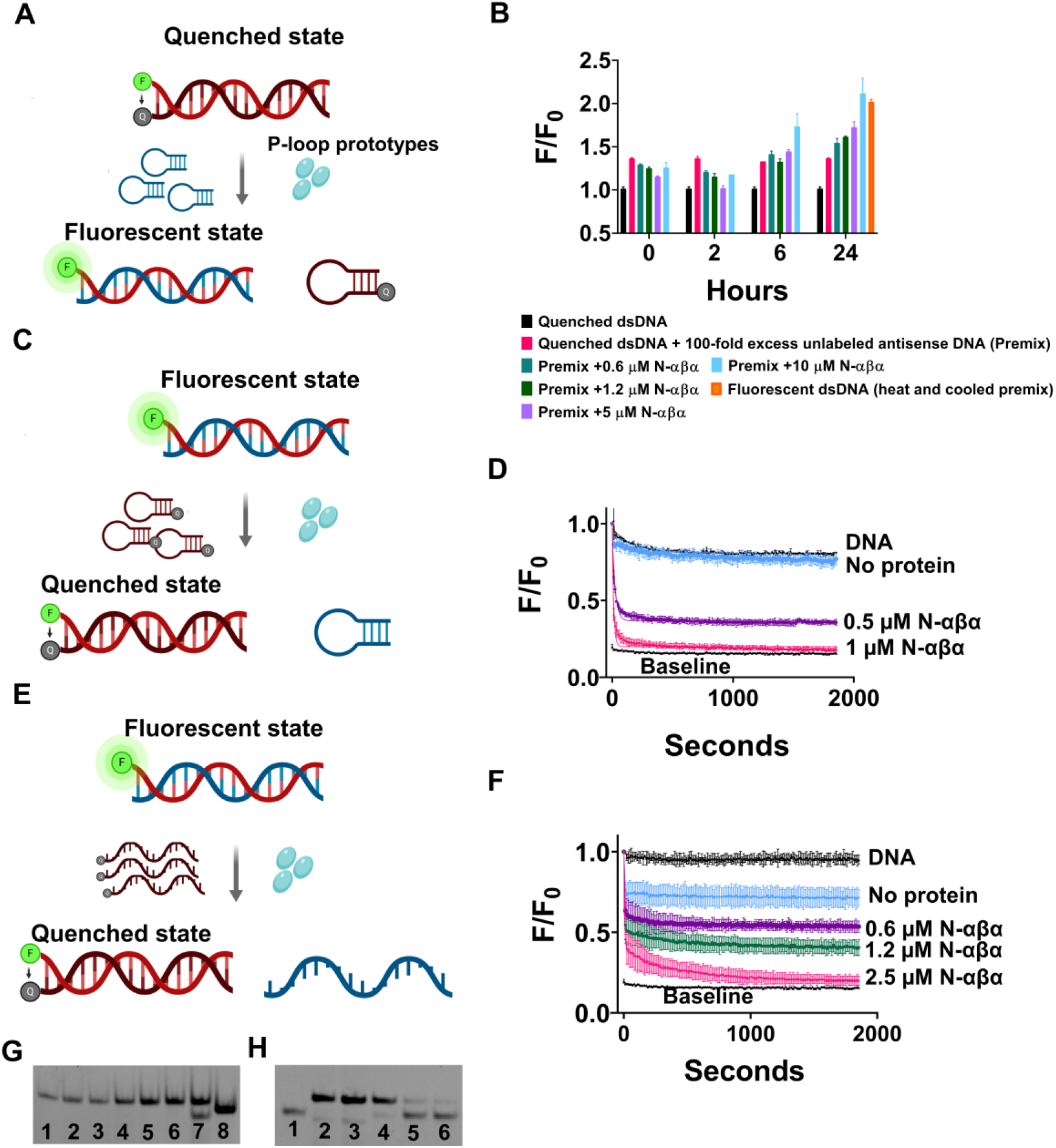
The N-αβα P-loop prototype mediates strand exchanges. **A.** A schematic of the strand exchange reaction between a quenched dsDNA and a non-labelled complementary strand (hairpin-forming). **B.** The exchange reaction was monitored by increase in fluorescence upon addition of N-αβα to a premix of quenched dsDNA and 100-fold excess of unlabeled complementary antisense strand. Fluorescence was normalized (F/F_0_) to quenched dsDNA (equals 1) and plotted vs. time (n = 2 to 4; error bars represent SD values). **C.** A schematic of the strand exchange reaction between a fluorescent dsDNA and a quencher containing complementary strand (hairpin forming). **D.** Fluorescence quenching (F/F_0_) was monitored upon addition of N-αβα to a premix of fluorescently labeled dsDNA and 10-fold excess of a hairpin-forming competing strand (hairpin forming) (n = 2 to 4; error bars represent SD values). Data were fit to a one-phase exponential decay (**Supplementary Table S8**). **E.** A schematic of the strand exchange reaction between a fluorescent dsDNA and a quencher containing complementary strand (linear). **F.** Fluorescence quenching (F/F_0_) was monitored upon addition of N-αβα to a premix of fluorescently labeled dsDNA and 10-fold excess of a linear competing strand (n = 2; error bars represent SD values). Data were fit to a two-phase exponential decay. **G.** Native EMSA gel of the strand-exchange reaction shown in panel *B*. The reactions were allowed to reach completion (24 hrs.) and DNA products were resolved on native polyacrylamide TBE gels (see Methods). The lanes are as follows: *1.* Quenched dsDNA (FAM-GA sense strand *plus* BHQ-1 antisense strand); *2.* DNA Premix (quenched dsDNA with 100-fold excess of unlabeled antisense strand); 3-6. DNA Premix with N-αβα (0.6, 1.2, 5 and 10 μM respectively); *7.* Fluorescent beacon dsDNA; *8.* Fluorescent ssDNA (FAM-GA sense strand) (**Supplementary Table S1**). Short dsDNA duplexes comprising of A/T rich regions are prone to improper annealing and/or formation of internal hairpins due to self-complementary regions. This is likely the reason for the visible lower band in lane 7. **H.** Native EMSA gel of a strand-exchange reaction as shown in panel *E* but with both strands fluorescently labeled. Strand-exchange reactions were carried out as described above, allowed to reach steady state (14 hr.), and analyzed on a native polyacrylamide TBE gel. The lanes are as follows: *1.* Fluorescent ssDNA (FAM-TC-linear antisense strand); *2*. Fluorescent dsDNA (FAM-GA-linear sense strand *plus* FAM-TC-linear antisense strand); *3.* DNA premix (fluorescent dsDNA with 10-fold excess of BHQ-1-linear antisense strand); *4.* DNA Premix with 0.6 μM N-αβα and *5.* with 1.25 μM N-αβα; *6.* Quenched dsDNA (FAM-GA-linear sense strand *plus* BHQ-1-linear antisense strand) with FAM-TC-linear antisense strand (5 nM; 1:1 ratio).

That the fluorophore-quencher interaction stabilizes the duplex DNA and hinders strand separation and exchange was indicated in the swapped setup showing rapid exchange – namely, displacement of an unlabeled strand by a quencher-containing strand (**Figure 3C**). In contrast to the set-up shown in **Figure 3A**, N-αβα mediated strand-exchange at the minute time scale, with only 10-fold excess of the complementary strand (**Figure 3D**; **Supplementary Table S8**). Acceleration of strand exchange by N-αβα was also also observed in the same setup yet with ‘linear’ ssDNA oligos that do not form a hairpin (**Figure 3E**, **3F**). Here too, no siginficant exchange occurred within the time scale of the experiment in the absence of N-αβα, while addition of ≤ 2.5 μM N-αβα induced complete exchange within minutes (**Figure 3D**, **3F**). Note that the concentration of N-αβα remains in large excess (≥ 0.5 μM) over the excess of competing ssDNA (50 nM). Overall, as expected, the energy gap between the start and end states dictates whether binding by the P-loop prototypes can shift the equilibrium between ds- and ssDNA, and also how fast the shift is. However, the N-αβα prototype also accelerated strand exchange in a more demanding setup (**Figure 3B**).

That the P-loop prototypes mediate strand-exchange was further validated by EMSA experiments. Once the strand-exchange reactions described above reached steady-state the bound proteins were removed by addition of hexametaphosphate and the DNA products were resolved on native polyacrylamide gels (**Figure 3G, 3H**). **Figure 3G** shows the transition from the initial quenched dsDNA to a fluorescent dsDNA, in protein concentration dependent manner, indicating DNA unwinding and exchange between the quencher labeled strand and an unlabeled complementary strand. However, this assay does not track down the change in state of the quencher-containing strand from ds-to ss-DNA. To this end, we used an alternate experimental set-up that follows **Figure 3E**, except that both strands of the duplex DNA were fluorescently labeled. Strand-exchange reactions with a complementary quencher-carrying strand were performed as described above and the DNA products were resolved on a native EMSA gel. The gel analysis (**Figure 3H)** indicated the transition of fluorescent dsDNA to a quenched dsDNA state (due to exchange of the bottom fluorescent strand with the quencher-containing strand) and a concomitant accumulation of the displaced flurescent strand (as indicated by the lower band in lanes 4 and 5). Overall, the observed changes in intensity of dsDNA bands (**Figure 3G, 3H**) and the parallel accumulation of ssDNA products (**Figure 3H**) occur only in the N-αβα containing reactions, consistent with the fluorescent assays, thus corroborating the helicase-like activity of P-loop prototypes.

### ssDNA release upon NTP and polyphosphate binding

Strand-separation as shown so far is, in essence, a shift in equilibrium in favor of P-loop prototype bound ssDNA. However, helicases are enzymes that turnover, namely they also release the bound DNA, typically upon ATP hydrolysis. The P-loop prototypes bind ssDNA and ATP via the same P-loop motif as suggested by mutations in the P-loop residues that abrogated binding of both^16^. Based on this observation, we asked if ATP addition could displace the ssDNA, thus also allowing the DNA to relax to its initial dsDNA state (**Figure 4A**). To this end, the N-αβα prototype was mixed with the beacon dsDNA and strand-separation was allowed to reach equilibrium. Upon subsequent addition of ATP or GTP, fluorescence reverted to its initial state indicating the release of the bound prototype and reversion to the starting dsDNA state (**Figure 4B**). Release by ATP occurred with an apparent K_D_ value in the range expected for a small ligand (K_D_^App^ = 2.8 mM) and GTP was a more potent inducer of reversion with 2-fold tighter K_D_^App^ (1.4 mM; **Figure 4C**). It seems, however, that release of the ssDNA is primarily conferred by the triphosphate group of these NTPs, and not the nucleoside moiety, as indicated by even tighter binding of triphosphate (K_D_^App^ = 790 μM). Given that inorganic phosphoanhydrides were proposed to have preceded NTPs as life’s energy coin, we tested polyphosphate – Kornberg’s energy fossil^35^ – as well as hexametaphosphate. N-αβα showed a strong preference for both polyphosphates (K_D_^App^ = 13.7 μM; calculated for an average of 18 phosphates per molecule, and 5.6 μM for hexametaphosphate; **Figure 4C**). Addition of phosphate at molar concentrations that are >100-fold higher than hexametaphosphate did not release the bound protein (5.6 mM phosphate barely had any effect on releasing the bound ssDNA, whereas 5.6 μM hexametaphosphate, equivalent to 33.6 μM phosphate, induced complete release; **Figure 4C**). Thus, the regain of fluorescence seems to be driven by specific ligand binding rather than by nonspecific effects such as changes in ionic strength. The efficient release of prototype-bound ssDNA by hexametaphosphate was further corroborated by ELISA experiments showing that prior incubation of N-αβα with hexametaphosphate inhibited binding to ssDNA (**Figure 4D**).

**Figure 4:**
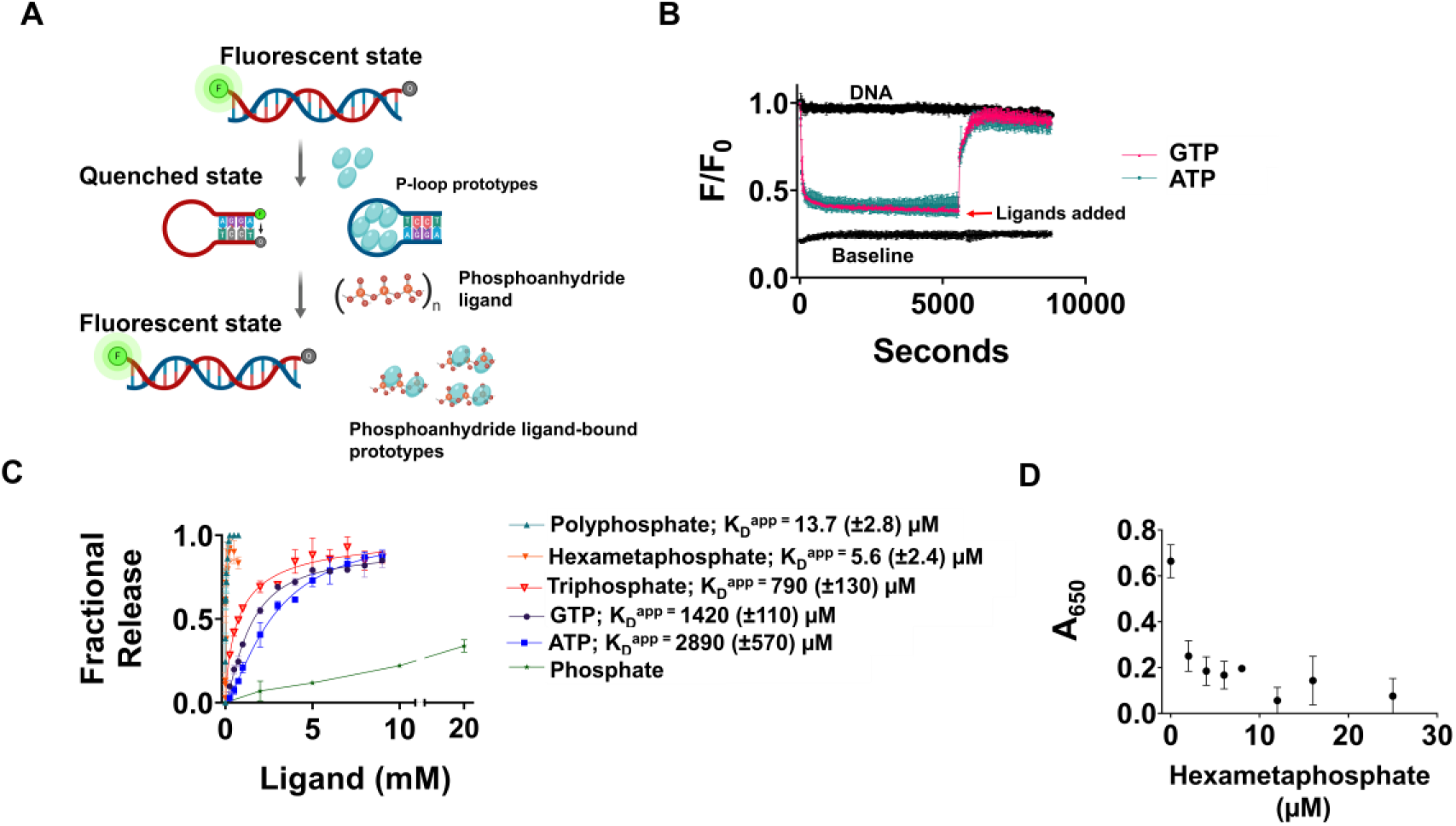
Release of bound ssDNA by phosphoranhydride ligands. **A.** A schematic description of the induction of strand separation by a P-loop prototype (1^st^ step) followed by the displacement of the bound ssDNA by various phosphoanhydrides (P⊓) and relaxation to the initial dsDNA state (2^nd^ step). **B.** Representative strand separation experiments with 0.4 μM N-αβα added at time zero. Subsequent addition of ATP (turquoise) and GTP (pink, both at 9 mM concentration) after 90 mins, leads to the release of the bound ssDNA and its relaxation to the initial dsDNA state. **C.** The apparent affinities (K_D_^App^) of binding of various phosphoanhydride ligands to P-loop prototype N-αβα. The fraction of released ssDNA upon addition of phospho-ligands was calculated by normalizing the F/F_0_ values from the plot in panel B. Complete release (=1) corresponds to the initial unbound dsDNA state and no release (=0) corresponds to the steady-state value of F/F_0_ prior to ligand addition. The apparent binding affinities (K_D_^app^) were calculated by Eq. 3 (**Supplementary Information**). Vertical error bars, and values in parenthesis, represent standard deviation from two independent experiments. **D.** Inhibition of ssDNA binding. Preincubation of 0.125 μM N-αβα with increasing concentrations of hexametaphosphate shows abrogation of binding to 24 base biotinylated ssDNA (**Supplementary Table S1**) in an ELISA-format as detected by anti-Histag antibodies (n = 2 to 4; error bars represent SD values).

### Quaternary structural plasticity

The intact P-loop prototypes tend to form dimers^16^. We assumed that their fragments are more likely to do so and possibly form even higher-order oligomers. Despite extensive attempts, we could not obtain crystals of N-αβα, or of any other functional construct described here. However, we performed chemical crosslinking experiments that revealed that N-αβα forms higher order oligomers, spanning from dimers to hexamers (**Supplementary Figure S2D**). Native mass spectrometry (MS) revealed that N-αβα self-assembles to form an ensemble of even higher-order forms, with the predominant and unambiguously assigned assemblies being 10-mer and 30-mer (**Figure 5A**). Furthermore, MS/MS experiments confirmed the identity of these species, as indicated, for example, by isolating the 30+ charge state of the 30-mer species, and inducing its dissociation into a highly charged monomer and a stripped 29-mer complex (**Figure 5B**). Dynamic light scattering (DLS) measurements also indicated that N-αβα exists in higher oligomeric form. The addition of inorganic polyphosphates to N-αβα results in an increase in the intensity of scattered light which translates to an increase in the average particle diameter (**Supplementary Figure S6**; **Supplementary Table S9**). In agreement with its lower apparent binding affinity (**Figure 4C**), GTP induced a smaller effect and the shift demanded higher concentrations. However, the diameter reported by DLS is only a proxy of the actual size, and the precise nature of the changes in the oligomeric assembly of N-αβα in response to ligand-binding needs to be further investigated.

**Figure 5.**
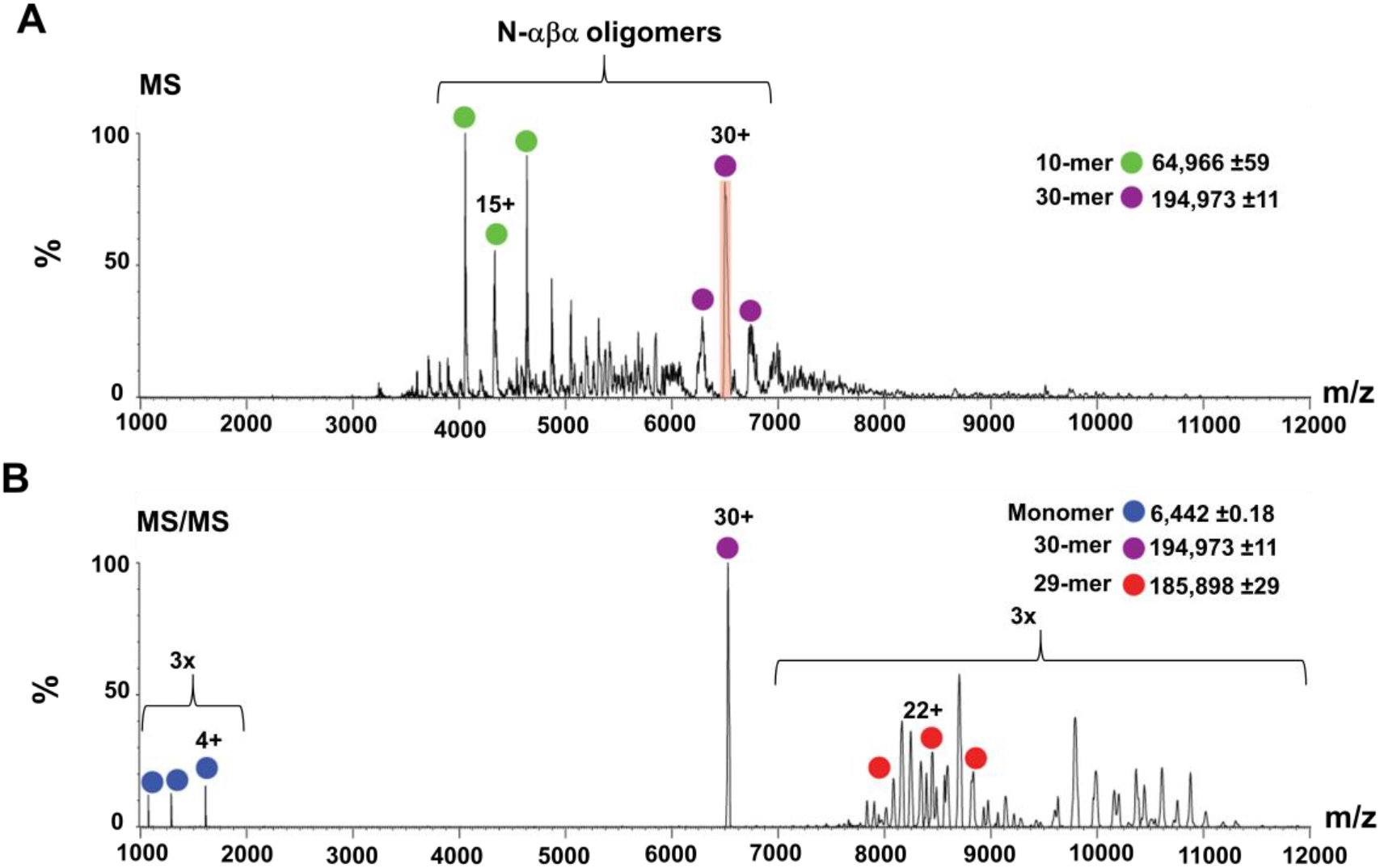
Quaternary structural characterization of N-αβα. **A.** Native MS analysis of N-αβα prototype. Under non-dissociative conditions, charge series corresponding to 10 and 30 N-αβα oligomers were unambiguously assigned in the MS spectrum (green and purple dots, respectively). The 30+ charge state that was selected for tandem MS analysis is shadowed in red. **B.** Tandem MS of the N-αβα 30-mer oligomer releases a highly charged monomer (blue; at the lower m/z range) and a stripped 29-mer oligomer (red), confirming the oligomer stoichiometry. The spectra are magnified 3-fold above 7,000 and below 2,000 *m/z*.

## Discussion

Previous studies have shown that NTP binding and modest catalysis can arise in polypeptides that comprise fragments of modern NTPases^16,36^ or of other ancient domains^37^. This study extends our previous report^16^ to demonstrate that polypeptides comprising the ancestral β-(P-loop)-α motif, with some minimal sequence additions to facilitate solubility, confer helicase-like functions. Our findings have several implications regarding the primordial P-loop NTPases, as discussed in the sections below.

### From a generalist phospho-ligand binder to specialized enzymes

In contemporary helicases, translocases and RecA proteins, the P-loop mediates NTP binding and hydrolysis, while DNA binding is mediated by other surface loops or even by a separate domain^20–23^. In contrast, in our prototypes, the P-loop binds NTPs and inorganic polyphosphates, as well as nucleic acids. This multifunctionality is the key to the prototype’s helicase-like action. Specifically, the strand-exchange assays demonstrate the ability of the P-loop prototypes to accelerate the exchange between a free and a duplex-bound DNA strand (**Figure 3**). However, in the unwinding experiments (**Figure 2**), binding of the P-loop prototypes merely shifts the equilibrium toward ssDNA. Nonetheless, the prototypes’ action can be reversed by addition of phospho-ligands such as ATP, and foremost by inorganic polyphosphates. These binding-release cycles are not enzymatic turnovers, yet may be a first step toward a *bona fide* helicase.

This multifunctionality is surprising, as although nucleic acids have phosphate groups, they fundamentally differ from NTPs. Can vestiges of a generalist P-loop, and specifically of ssRNA/DNA binding, be found in extant P-loop NTPases? We searched the PDB for domains that belong to the P-loop NTPase lineage and have ssRNA/DNA bound in proximity to the P-loop (see Methods). This search identified at least two extant P-loop NTPase families: XPD helicase (**Figure 6A**), and polynucleotide kinase (**Figure 6B**), where the P-loop interacts with ssDNA, as it does in our P-loop prototypes. In the case of XPD helicase, the canonical Walker A motif diverged and phosphate binding is mediated by a short motif, SGR, at the tip of α1. Remarkably, in case of polynucleotide kinase (**Figure 6B**), a canonical P-loop binds GTP/GDP as well as ssDNA (oligonucleotide) by a network of interactions facilitated by the P-loop residues. This, to our knowledge, is the only instance of an extant multifunctional P-loop that not only catalyzes phosphoryl transfer, but also binds ssDNA.

**Figure 6.**
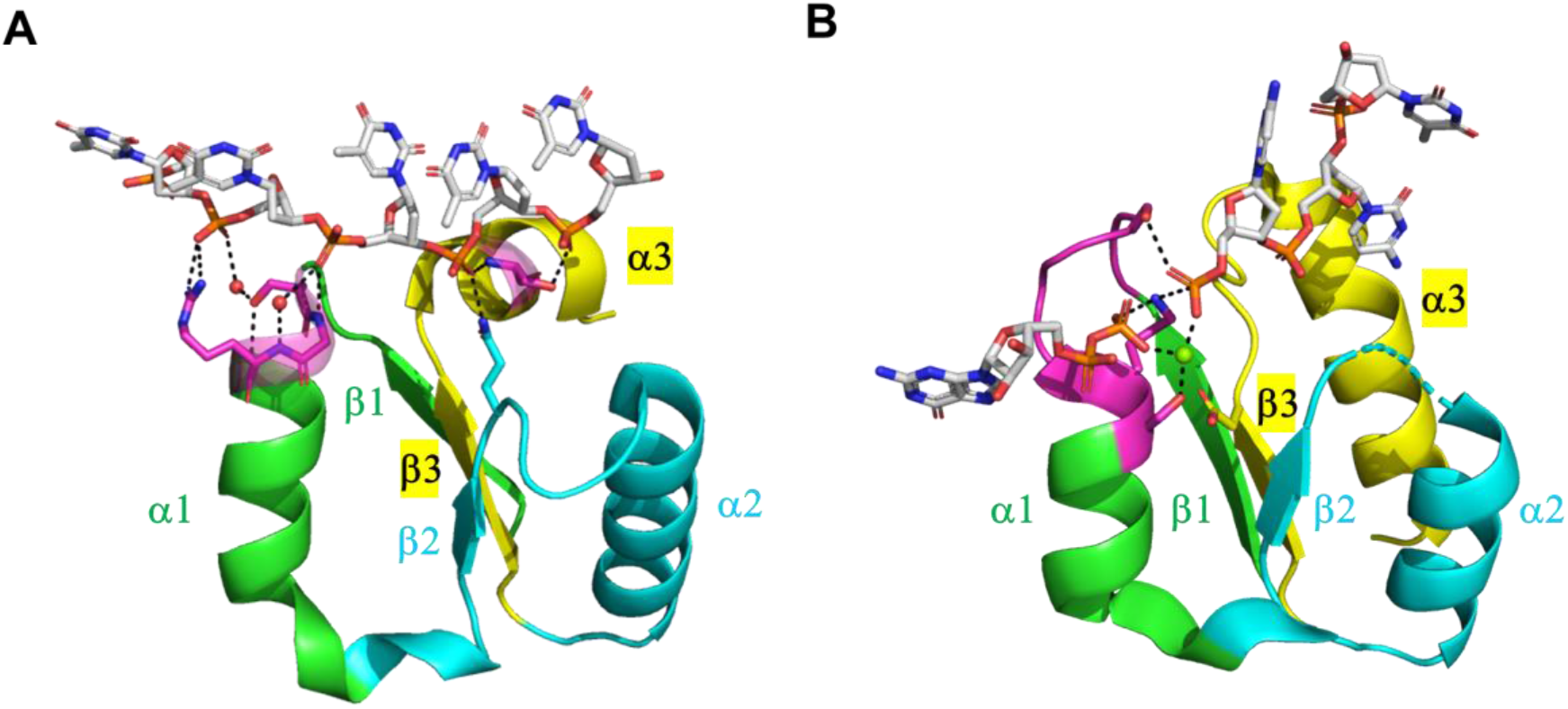
Instances of P-loops binding to ssDNA in extant P-loop NTPases. **A.** The Helicase_C_2 domain of XPD helicases (F-groups, ECOD: 2004.1.1.106, Pfam: PF13307). Shown here is a fragment taken from a representative structure (ATP dependent DinG helicase; ECOD domain ID: e6fwrA1, residues 448-703; the fragment shown, residues 535-599, spans from β1 to α3). The strand topology of this domain follows the simplest P-loop NTPase topology (2-3-1-4-5). Its P-loop resides at the tip of α1, in the very same location as the Walker A, but its sequence is non-canonical (SGR, in magenta). Shown are direct as well as water-mediated interactions between the residues of the P-loop and phosphate groups of the ssDNA oligonucleotide (waters shown as red spheres). A glutamine at the tip of β2 (in cyan), and a serine at the N-terminus of α3 (in magenta) provide additional anchoring points for the ssDNA. **B.** Bacterial polynucleotide kinase (F-groups, ECOD: 2004.1.1.32; Pfam: PF13671; strand topology: 2-3-1-4-5). Shown here is a fragment of the P-loop NTPase domain from a representative structure *(Clostridium thermocellum* polynucleotide kinase; ECOD domain ID: e4mdeA1, residues 1-170; the shown fragment, residues 8-37 and 62-98, spans from β1 to α3, with residues 38-61 truncated for clarity). The canonical Walker A motif resides between β1 and α1 (in magenta, GSSGSGKS) and binds GTP. This representative structure shows the product complex of GDPoMg^+2^ and the phosphorylated ssDNA. The lysine and serine residues of the P-loop motif at the tip of α1 provide bridging interactions, coordinated by an Mg^2+^ ion (green sphere), between the β phosphate of the bound GDP and the phosphate group at the 5’-OH of the ssDNA.

At present, in the absence of structures at atomic resolution, the precise mode of binding by our P-loop prototypes remains unknown. It seems, however, that the Walker A motif is not critical perse. Specifically, we observed that the first Gly of the Walker A motif of our P-loop prototypes plays no role, and that the last three residues (GK(S/T)) seem most critical; however, even at these three positions, mutations reduce but not completely abolish binding (see Ref.^16^ and **Supplementary Figure S3**). Indeed, the most rudimentary forms of phosphate binding make use of backbone amides, of glycine as well as other residues, and most critically, of bidentate interactions involving both backbone and side-chain H-bonds, foremost by Ser/Thr^14^. These elements are seen in XPD helicase (**Figure 6A**) and are likely to be the key to a generalist P-loop.

That the ancient P-loop was a ‘generalist’ that mediates multiple functions is in line with the hypothesis that the earliest enzymes were multifunctional^38^. A plausible explanation, in accord with the Dayhoff’s hypothesis^3,4^, is that such multi-functional primordial P-loop fragments were, at a later stage, duplicated and fused. This resulted in a tandem repeat of β-(P-loop)-α elements, where the P-loops comprise the active site. Further divergence allowed one P-loop to retain NTP binding and develop the ability to catalyze phosphoryl transfer, while the other P-loop(s) were repurposed to bind nucleic acids. This scenario is illustrated by XPD helicases wherein a P-loop ATPase domain, the DEAD domain (ECOD domain ID: e6fwrA2), is fused to the ssDNA binding helicase C_2 domain.

### A minimal structural context for P-loop function

Bioinformatics analyses have pointed out the β-(P-loop)-α-β fragment as the recurrent structural/functional unit in dinucleotide binding proteins^5,39^. While we found that this fragment mediates strand separation, it was aggregation-prone, likely due to the highly hydrophobic βl strand (**Supplementary Table S3**), and active only under non-stringent conditions. In contrast, the α-β-(P-loop)-α fragment was soluble and functional in the presence of salt. Having a helix preceding the β-(P-loop)-α fragment seems to present an advantage, primarily by enhancing solubility of this exceedingly hydrophobic βα fragment. We also note that although in most P-loop NTPase domains β1, from which the P-loop extends, comprises the N-terminus, an alpha helix that precedes β1 is also in some P-loop families, *e.g.* in the F-type ATP synthases (F-groups, ECOD: 2004.1.1.53; Pfam: PF00006), Transcription termination factor Rho (F-groups, ECOD: 2004.1.1.53; Pfam: PF00006), and RecA (F-groups, ECOD: 2004.1.1.237; Pfam: PF00154). Moreover, the helicase-like function is retained despite truncation of most of this N-terminal helix, and even the β-(P-loop)-α fragment that could not be isolated was co-purified with bound nucleic acids. These results, along with the observation that ancestral β-(P-loop)-α motif can be embedded in various structural contexts while retaining function, suggest that β-(P-loop)-α is likely the minimal stand-alone seed from which P-loop NTPases could have emerged.

### Dynamic self-assembly is key to function

We hypothesized that self-assembly is a critical bridging step that enables short polypeptides to be functional on their own prior to their duplication and fusion—a step that likely required advanced genetic and protein translation machineries^4^. Shorter polypeptides are unlikely to confer biochemical functions such as small ligands binding^40^. However, self-assembly, even in rudimentary forms such as amyloid fibers^41,42^ or coacervates^43^, can provide the operative volume and network of interactions necessary for biochemical function^42,44^. Although further studies are required to elucidate the mechanistic and structural basis of these assemblies, the propensity of the P-loop prototypes to self-assemble is evident (**Figure 5**). Foremost, oligomerization confers avidity, *i.e.*, binding via multiple P-loops per functional unit. Otherwise, binding at μM affinity by a solvent exposed loop would have been impossible. Changes between oligomeric states are also indicated by the cooperativity of the strand separation reactions, as manifested in high Hill coefficients and complex kinetics (**Figure 2**). Similar DNA-induced cooperative functions are documented for SSB proteins^45^, helicases^46,47^ and RecA proteins^48–51^, and the latter two are also P-loop NTPases. Further, RecA has been shown to also possess strand-separation activity on short dsDNAs^52^, and its recombinase function involves changes in oligomeric states, from monomers-dimers up to long filaments^1,2^.

### The potential role of P-loop prototypes in the pre-LUCA world

Whether proteins (metabolism) or nucleic acids (information) came first has been extensively debated. However, these scenarios are not necessarily mutually exclusive – both elements had to coexist in some rudimentary form well before the LUCA^53^. It is widely accepted that P-loop NTPases emerged early, possibly within an RNA-protein world^5,11–13,17,54,55^. Accordingly, RNA/DNA remodelers such as SFII helicases^55^ and RecA constitute a major class of P-loop NTPases ascribed to the LUCA^7,8^. P-loop polypeptides may have therefore marked the early stages of cooperation between proteins and nucleic acids (other examples include polypeptides that interacted with ribosomal RNA^56–58^). Specifically, in the absence of protein-based polymerases, nucleic acids must have replicated either non-enzymatically^59,60^ and/or in a self-catalytic manner^61^. However, in such scenarios, a fundamental impediment for multiple rounds of replication is the stability of duplex RNA^62^ (or DNA for that matter^63^). Heat can separate the strands, yet reannealing would be faster than any abiotic replication^64^. A plausible solution for unwinding RNA duplex products is short “invader” oligonucleotides^65^; however, how strand-displacement would initiate remains unresolved^65^. Thus, for multi-round replications, certainly of templates that are dozens of nucleotides long, an unwinding polypeptide is likely necessary. The duplex unwinding and strand displacement activities of our P-loop prototypes provide a plausible solution to these challenges.

Our results suggest that the strand-separation and exchange mediated by the P-loop prototypes is a thermodynamically driven process. The feasibility and rate of this process are dictated by the energetic gap between the initial and final DNA states. It is therefore likely that prior to the emergence of enzymatic function, primordial helicase-like polypeptides were fundamentaly ssDNA/RNA binding proteins that ‘passively’ unwound DNA duplexes by relying on transient exposure of ssDNA due to thermal fluctuations or/and DNA fraying^27,28^. Considering that the prototypes’ affinity to the GA-rich strand of the beacon is low, it is likely that strand separation is primarily driven by binding to the TC-rich strand (**Figure 2F**). In fact, this is a commonly reported mode of unwinding for hexameric replicative helicases where the helicase-ring encircles one strand while the other strand is excluded^66^. The ability to bind ssDNA likely undelies the strandexchange process as well (**Figure 3**) thus resembling the mode of action of extant RecA proteins^67^. Thus, although further studies will be required to decipher the precise mechanism of action of these P-loop prototypes, it appears that these rudimentary P-loop helicases hold tangible links to their modern descendants.

### Inorganic polyphosphates as energy fossils

Our P-loop prototypes bind phosphate moieties with no apparent interactions with the nucleoside (ATP, GTP and triphosphate bind similarly; **Figure 4**). Foremost, long chain polyphosphate binds with 1000-fold higher affinity than ATP. Indeed, inorganic phosphoanhydrides were proposed to have been the abiotic energy precursors of NTPs^68^. Of particular interest is hexametaphosphate – a cyclic ring of six phosphates that also binds with μM affinity (**Figure 4**). In the primitive biotic world, polyphosphates could have served not only as an energy source, but also as a scaffold that facilitated the assembly and orientation of phospholipids, nucleic acids and proteins^69^. Further, trimetaphosphate has been extensively explored as an abiotic condensation reagent that promotes synthesis of peptides and nucleic acids^70^. Trimetaphosphate binds weakly to our P-loop prototypes, but it coexists in equilibrium with hexametaphosphate^70^ that binds at μM affinity. Thus, the mode of action of these P-loop prototypes is tantalizingly tailored to the requirements of a primordial world.

**To conclude**, Charles Darwin’s immortal statement echoes: “from so simple a beginning endless forms most beautiful and most wonderful have been, and are being, evolved”. The P-loop prototypes described here provide a glimpse of how these simple beginnings might have looked – short and simple polypeptides that are nonetheless linked to their modern descents in sequence, structure and function.

## Materials and Methods

P-loop prototypes were cloned, expressed and purified as described^16^ with some modifications (**Supplementary text: Materials and Methods**). For strand-separation assays, fluorescence quenching was measured upon titrating the beacon dsDNA with varying concentrations (0.1 to 1 μM) of P-loop prototypes in 50 mM Tris pH 8, or in 50 mM Tris with 100 mM NaCl (pH 8). Reactions were monitored for two hours using the Infinite M Plex microplate reader (TECAN), at 24 °C, with an excitation/emission wavelength of 495/540 nm. The fluorescence decay values were normalized to the initial fluorescence (before protein addition; F0) to derive F/F_0_ values that were fit to standard one-or two-phase exponential decay equations (**Eq. 1 and 2; Supplementary text: Materials and Methods**) to derive the apparent rate constants (kapp). The apparent binding affinities (K_D_^app^) and Hill’s coefficient (h) (for strand-separation and release by phospho-ligands) were determined by plotting the normalized end-point F/F_0_ values (denoted as ‘Fraction Unwound’) against protein concentration and by fitting the data to a cooperative binding equation (**Eq. 3, Supplementary text: Materials and Methods**).

Strand-exchange assays were performed by titrating DNA premixes (dsDNA with excess of complementary single strand) (**Supplementary Table S1**) with varying concentrations of the N-αβα prototype (0.5 to 10 μM). The reactions were carried out in the stringent binding conditions (50 mM Tris with 100 mM NaCl; pH 8) and monitored using the Infinite M Plex microplate reader for two hours as above. The normalized end-point F/F_0_ values were fit to standard one or two exponential decay equation (**Eq. 1 and 2**).

For fluorescence anisotropy measurements, various ss and dsDNA constructs (**Supplementary Table S1**) were titrated with N-αβα prototype (0.1 to 2 μM) in buffer containing 50 mM Tris with 100 mM NaCl (pH 8) at 24 °C. Fluorescence polarization was monitored for two hours using the Cytation 5 multi-mode reader (BioTek) with the green filter set (485/20 nm for excitation, and 528/20 nM for emission) and a top optic probe with a 510 nm dichroic mirror. The steady-state anisotropy values were derived from the parallel and perpendicular emission intensities using the standard anisotropy equation (**Eq. 4, Supplementary text: Materials and Methods**), and binding affinities (K_D_) and Hill’s coefficient (h) were derived by fitting to a cooperative binding equation (**Eq. 5, Supplementary text: Materials and Methods**). For anisotropy-quenching dual assays, the change in anisotropy and fraction unwound were measured as described above and plotted against protein concentration. All data fitting was carried out using GraphPad Prism (8.3.0) software.

For native mass-spectrometry measurements, the N-αβα polypeptide was dialyzed against 150 mM ammonium acetate (pH 7.5) and diluted to a final concertation of 10 μM. Nanoflow electrospray ionization MS and tandem MS experiments were conducted under non-denaturing conditions on a Q Exactive UHMR Hybrid Quadrupole-Orbitrap mass spectrometer (Thermo Fisher Scientific).

Identification of ssDNA binding by extant P-loop NTPases was carried out using a short script implemented in Python to identify structures, from the Protein Data Bank, in which a P-loop domain and ssDNA come within 3.5 Å of each other. *(For more details, see **Supplementary text: Materials and Methods***)

## Acknowledgments

This research is funded by a Minerva Foundation grant for scientific cooperation between Germany and Israel. We are grateful to Prof. Ita Gruic-Sovulj for her insightful suggestion to simultaneously monitor quenching and anisotropy. We thank Dr. Guy Shmul for help with DLS analysis and for providing size standards and Dr. Gili Ben-Nissan for help with native massspectrometry experiments. Schematics for strand-separation, strand exchange and release assays were created with BioRender.com.

