## Supplementary Information file for "Helicase-Like Functions in Phosphate Loop Containing Beta-Alpha Polypeptides"

**for**

**This PDF file includes:**

Supplementary text: Materials and Methods

Figures S1 to S6

Tables S1 to S9

References for supplementary information

**Supplementary text: Materials and Methods**

**Protein expression and purification**

Synthetic gene fragments encoding P-loop prototypes were purchased from *Twist Bioscience*. Cloning of synthetic genes and generation of mutant constructs, were performed as described previously ^1^. All constructs had a C-terminal tag, comprising a Trp reside for concentration determination by measuring absorbance at 280 nm, (the P-loop prototypes have no aromatic residues) followed by 6xHis for purification (DNA and amino acid sequences are provided in **Supplementary Tables S2 and S3**). All proteins were expressed in *Escherichia coli* BL21 (DE3) cells. Cells were grown in LB media at 37 ºC with shaking (220 rpm) until OD600 ~0.6 and protein expression was induced by the addition of ITPG (1 mM final concentration) at 20 ºC. Following overnight induction at 20 ºC with shaking, cells were harvested by centrifugation (4000 g for 15 min at 4ºC) and resuspended in lysis buffer: 50 mM sodium phosphate, 20 mM imidazole, 500 mM NaCl, 6 M guanidine hydrochloride (pH 7.4), supplemented with the EDTA-free antiprotease cocktail supplied by *Sigma*. Cells were lysed by sonication, the debris was centrifuged down for 30 mins at 13000 rpm, and the supernatant was filtered through 0.22 µm membrane filter (*Merck Millipore*). Clarified lysates were loaded onto a HisTrapTM FF 5 mL column (*GE Healthcare*) using an AKTA FPLC. The guanidine hydrochloride that was added to dissociate bound nucleic acids was removed using a linear gradient of 12 column volumes of 50 mM sodium phosphate, 20 mM imidazole, 500 mM NaCl, pH 7.4. Bound proteins were eluted using one-step elution with 50 mM sodium phosphate, 500 mM imidazole, 500 mM NaCl, pH 7.4. Protein yield and purity was assessed by SDS-PAGE. Two rounds of dialysis were carried out to dialyze out the imidazole. The P-loop prototypes generally precipitated during the dialysis to remove imidazole. Precipitated protein samples were centrifuged, the pellet was collected and dissolved in buffer containing 50 mM Tris, 150 mM NaCl and 1 M L-arginine (pH 8). Purified prototypes stored in this buffer at 4 °C, at 100 to 200 µM concentrations, remained mostly soluble and active for periods of seven to ten days.

**Strand separation and exchange assays**

Synthetic DNA oligonucleotides for unwinding and strand-exchange assays were obtained from *Integrated DNA Technologies, Inc*, and *Metabion International AG*. Their sequences, and which experiments they were used, are listed in **Supplementary Table S1**. The beacon dsDNA was prepared by hybridizing the beacon oligo (GA-beacon sense strand) and the unlabeled antisense oligo each at 10 µM in 50 mM Tris (pH 8). Hybridization was done by heating the oligo mixture at 95 °C for 5 mins followed by slow cooling overnight. Strand separation reactions were performed in a buffer containing 50 mM Tris (pH 8), and at the stringent binding conditions in 50 mM Tris (pH 8) with 100 mM NaCl. The reaction was initiated by adding P-loop prototypes (1 to 0.1 µM) to beacon dsDNA (5 nM) in 100 µl reaction volumes, at 24 °C, in Nunc-flat black 96-well plates. Fluorescence was monitored for two hours using Infinite M Plex microplate reader (*Tecan*) with an excitation/emission wavelength of 495/540 nm. Fluorescence decay values were normalized to the initial fluorescence before protein addition (F_0_) to obtain F/F_0_ values. The non-specific quenching values observed with 5 nM of beacon DNA digested overnight with Benzonase (*Merck Millipore*) was subtracted from the protein-derived signals. Rate constants were determined by fitting the time dependent F/F_0_ values to one-phase (Eq. 1) using the GraphPad Prism (8.3.0) software.

(1) F/F_0 (t)_ = (F/F_0_)^∞^ + (1 – (F/F_0_)^∞^) × *e* ^–kapp*t^

Where t is time, (F/F_0_)^∞^ is the end-point value of F/F_0_, k_app_ is the apparent first-order rate constant.

Strand-separation reactions with intact P-loop prototype (**main text; Figure 1B**) comprised of an initial fast phase followed by a slow phase. Rate constants were derived by fitting data to the biphasic exponential decay equation (Eq. 2) using the GraphPad Prism (8.3.0) software:

(2) F/F_0_ _(t)_ = (F/F_0_) ^∞^ + (1 – (F/F_0_) ^∞^) [A^fast^ × e^-(k^_app1_^*t)^ + ((1-Afast) × e^-(k^_app2_^*t))^]

Where t is time, (F/F_0_)^∞^ is the end-point value of F/F_0_, A^fast^ is the amplitude of fluorescence decay of the fast phase, k_app1_ and k_app2_ are the apparent first order rate constants for fast and slow phases respectively.

For determining apparent binding affinities (**main text; Figure 2C**), the end-point F/F_0_ values (two hours) from the strand-separation experiments were derived for each protein concentration (P), and normalized (0 = the starting, fluorescent beacon substrate; 1 = the assay baseline, i.e., the fully quenched hairpin state). Normalized F/F_0_ values (‘Fraction Unwound’) were then plotted against protein concentration. The resulting data were fitted to a binding isotherm equation (Eq. 3; GraphPad Prism) with cooperativity, to derive the apparent dissociation constant (K_D_^App^) and the Hill’s coefficient (h).

(3) F/F_0 (P)_ = [P]^h^/(K_D_^App^ + [P]^h^)

Strand exchange assays (**main text; Figure 3**) were carried out in 50 mM Tris (pH 8) with 100 mM NaCl. Stock solutions of dsDNA were prepared by hybridizing the FAM sense strand (FAM-GA sense strand or FAM-GA linear sense strand; **Supplementary Table S1**) with the BHQ-1 labeled or unlabeled TC-rich antisense strands (hairpin forming or linear; **Supplementary Table S1**) in 50 mM Tris and 100 mM NaCl (pH 8), at 10 µM each, by heating and slow cooling as above. A mixture of this hybridized dsDNA (5 nM) and BHQ-1 or unlabeled antisense strands (50 nM) (hairpin forming or linear; **Supplementary Table S1**) was then used as the DNA substrate and reactions were initiated by adding P-loop prototypes (0.5 to 10 μM). Changes in fluorescence were normalized to the initial state (prior to protein addition) and data were fit to Eq. 1 or 2 as described above.

For determining apparent binding affinities of phospho-ligands, the fraction of released ssDNA (**main text; Figure 4**), for each ligand concentration (P), was normalized (1 = complete release and reversion of ssDNA to the starting, fluorescent beacon dsDNA substrate; 0 = no release, i.e., without added ligand). Normalized F/F_0_ values were then plotted against the ligand concentration and the resulting data were fitted to a binding isotherm equation with cooperativity as described above (Eq. 3), to derive the apparent dissociation constant (K_D_^App^) and the Hill’s coefficient (h).

**Gel based demonstration of helicase-like activity**

Once the strand-exchange reactions described above have reached steady state (24 hr for **Figure 3G** and 14 hr for **Figure 3H**), 750-µM hexametaphosphate (*Sigma 305553*) was added to release the bound proteins. For experiment shown in Figure 3H, we observed displaced ssDNA products after the usual two-hour reaction time, however their clear visualization required the reactions to be incubated overnight (**Figure 3H**). This is likely because deproteinization of DNA samples prior to electrophoresis can introduce artifacts such as loss of displaced ssDNA along with the protein ^2^, The samples were immediately mixed with loading dye and the DNA products were loaded onto 10% (29:1 acrylamide:bisacrylamide) Tris-Borate EDTA (TBE). The gels were run at 160 V for 90 min, at room temperature, in 1xTBE buffer and scanned using Amersham Typhoon scanner using the 488/525 nm laser-filter set at 500 volt PMT (photo multiplier tube) setting.

**Fluorescence anisotropy measurements**

Fluorescence anisotropy measurements (**main text; Figure 2, Supplementary Figure S4C, S4D**) were carried out using Cytation 5 (BioTek) instrument at 24 °C. Here, 5 nM of DNA (6-FAM labeled, or dually labeled (6-FAM and BHQ-1), single or double strand DNA constructs, as listed in **Supplementary Table S1**) were titrated with varying concentrations of N-αβα (0.1 to 2 µM) in 100 µL reactions. Fluorescence polarization was measured using the green filter set (485/20 nm for excitation, and 528/20 nM for emission) and top optic probe with a 510 nm dichroic mirror. The reactions were monitored for two hours, and the data processed with the Gen5.0 (BioTek) software. The resultant parallel (Ivv) and perpendicular (Ivh) emission intensity values were then used to calculate the samples’ anisotropy with the standard equation (Eq. 4):

(4) r_(P)_ = (I_vv_ – (G * I_vh_))/(I_vv_ + (2 * G * I_vh_))

Where r(P) is the steady-state anisotropy (after two hrs incubation) at a given protein concentration, I_vv_ and I_vh_ are the parallel and perpendicular emission intensities, respectively, and G is the grating factor that corrects for instrumental variation in detection of vertically and horizontally polarized light. G is calculated as I_hv_/I_hh_; where I_hv_ is the emission intensity when the excitation polarizer is in the horizontal position and the emission polarizer in vertical position. I_hh_ is the emission intensity when both the excitation and the emission polarizers are in horizontal position.

The endpoint anisotropy values as calculated from eq. 4 were normalized to the anisotropy of the free DNA (the latter being equal to 0). The normalized values were then fitted to a binding isotherm equation with cooperativity (Eq. 5; GraphPad Prism), to derive the dissociation constant (K_D_) and Hill coefficient (h).

(5) r_(P)_ = r_max_ _*_ [P]^h^ /(K_D_ + [P]^h^)

r_max_ was set to a constant value of 0.2 and corresponds to the maximum observed anisotropy in our experimental setup; K_D_ is dissociation constant; h is Hill’s coefficient.

Fluorescence anisotropy-quenching dual assays (**main text; Figures 2D, 2E**) were performed using the same instrument and reaction conditions described above. For each protein concentration, the absolute level of fluorescence, and fluorescence polarization, were measured for a duration of two hours. The unwound fraction of DNA (‘Fraction Unwound’) and change in anisotropy were calculated as described above, and then applied to equations 3 and 5, respectively, using GraphPad Prism software as described above.

**Enzyme-linked immunosorbent assay (ELISA)**

ELISA experiments were performed as previously described ^1^ with some modifications. Stock solutions of 24 bp biotinylated dsDNA were prepared by hybridizing (see above) equimolar concentrations (100 µM) of biotinylated-24nt-ssDNA with the unlabeled (antisense) strand in buffer containing 50 mM sodium phosphate and 100 mM NaCl; pH 7 (oligo sequences are listed in **Supplementary Table S1**). Streptavidin coated plates (*StreptaWell, Roche*) were coated with 100 µl of 0.05 µM of biotinylated 24 bp dsDNA, or biotinylated-24nt-ssDNA, diluted in ELISA buffer (50 mM sodium phosphate and 100 mM NaCl; pH 7 and 1 % non-ionic detergent Nonidet P40 and 0.1% BSA, pH 7) for 1 hr at ambient temperature. In parallel, wells for background subtraction were left uncoated (incubated with 100 µl ELISA buffer). Following coating, wells were washed with ELISA buffer and 100 µl of P-loop prototypes solutions were added (at 0.25 to 0.06 µM) to both DNA-coated and background wells, and incubated for 1 hr. The unbound proteins were washed extensively with ELISA buffer (10 times) and 100 µl of 0.1 µg/ml HRP labeled mouse anti-His antibody (200 µg/ml, Santa Cruz Biotechnology) were added and incubated for 40 mins. The wells were washed extensively, 100 µl of substrate solution containing 1.25 mMol/L TMB and 2.21 mMol/L hydrogen peroxide, <1% dimethyl sulfoxide in 0.08 Mol/L acetate buffer at pH 4.9 (3, 3’, 5, 5’-tetramethybenzidine; TMB; ES001, Millipore) was added and absorbance at 650 nm was monitored for 30 mins. The results are presented as endpoint values of OD^650nm^ (A_650_) after subtraction of the background signal.

**Chemical crosslinking**

Chemical crosslinking of P-loop prototypes was performed with EDC (1-ethyl-3-(3-dimethylaminopropyl) carbodiimide, hydrochloride salt) ^3^. Reactions were carried out as follows: purified P-loop prototypes were dialyzed against 10 mM MES buffer (pH 4.5), and diluted to 30 µM concentration in the same buffer. In 20 µl reaction volumes, 30 µM of protein were mixed with EDC (10 to 60 mM final concentration) dissolved in the above buffer. Reactions were incubated for 30 mins at room temperature and quenched with adding 50-fold molar excess (over cross-linker concentration) of Tris buffer (pH 8). Samples were mixed with SDS loading dye and analyzed on 16% SDS-PAGE gels run with Tris-glycine buffer.

**Dynamic light scattering (DLS)**

DLS analysis of N-αβα construct was performed using Malvern Zetasizer Nano-ZS particle analyzer (*Malvern Panalytical*). The P-loop prototypes were diluted to 1 µM in buffer containing 50 mM Tris with 100 mM NaCl (pH 8). Following an equilibration time of 2 min, light scattering was monitored in a disposable cuvette in 1 ml volumes at 25 °C. The measurement duration was set to ‘automatic’, resulting in 11 to 17 runs for each measurement and three measurements were performed for each sample. The data were fit to a correlation function and analyzed by the method of cumulants ^4^ to obtain the Z-average diameter (nm) (i.e. mean particle size). To test the effect of phospho-ligands, the prototype was mixed with different phosphate ligands, and light scattering was monitored as described above to obtain the Z-average diameter (nm).

**Native mass spectrometry analysis**

Nanoflow electrospray ionization MS and tandem MS experiments (**main text; Figure 5**) were conducted under nondenaturing conditions on a Q Exactive UHMR Hybrid Quadrupole-Orbitrap mass spectrometer (Thermo Fisher Scientific). Prior to MS analysis, the purified N-αβα was dialyzed against 150 mM ammonium acetate (pH 7.5) and diluted to a final concertation of 10 µM. Spectra were collected under positive ion mode, and conditions were optimized to enable the ionization and removal of adducts while retaining the noncovalent interactions of the proteins tested. In MS/MS experiments, the relevant m/z values were isolated and argon gas was admitted to the collision cell. Typically, aliquots of 2 μL of sample were electrosprayed from gold-coated borosilicate capillaries prepared in-house ^5^. Spectra are shown using UniDec ^6^ for mass detection, and without smoothing or background subtraction.

The following experimental parameters were used on the UHMR Hybrid Quadrupole-Orbitrap platform: spray voltage 1.35 kV, inlet capillary temperature 250 °C. The trapping gas pressure was set to 5 corresponding to HV pressures of 1.5x10-4 mbar. To facilitate efficient desolvation of proteins, in source trapping was set to -100 V and high-energy collision dissociation was set to -10 eV. Mass spectra were recorded at a resolution of 3,125. Injection flatapole DC bias was set to 5 V. For tandem MS the precursor (6,540 m/z) was isolated and activated using the HCD cell at 100 eV. Instruments were externally mass-calibrated using a cesium iodide solution at a concentration of 2 mg/mL.

**Identification of ssDNA binding by P-loop NTPases**

All structures annotated by the ECOD database (version develop275) as containing a P-Loop domain (X-group 2004) were downloaded from the Protein Data Bank – 5,818 structures in total. Structures in which a P-Loop domain and ssDNA come within 3.5 Å of each other were identified using a short script implemented in Python. Hits were validated by visual inspection (**main text; Figure 6**).

**Supplementary Figures**
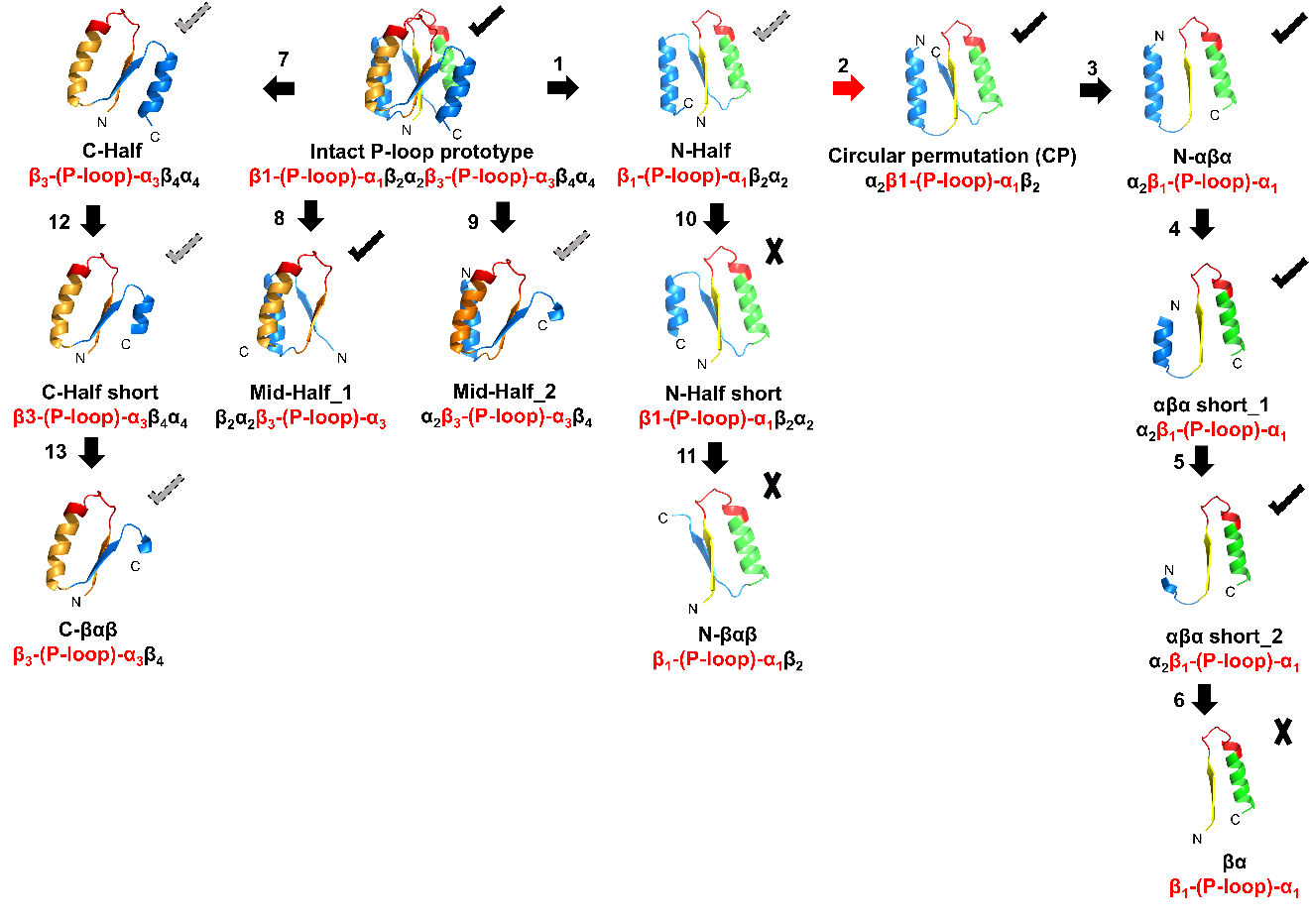
**Figure S1: Schematic representation of new P-loop prototypes used in this study.**

P-loop prototype fragments were constructed from the intact 110 residues prototype (D-Ploop; Ref ^1^.) by its truncation, or by circular permutation coupled with truncation. The fragments had either a ‘βαβα-type’ or ‘αβαβ-type’ architecture; specifically:

- Steps 1 to 6 began with the N-terminal half (N-half), and are described in main text (**Figure 1C**).
- Steps 7, 8 and 9 relate to the intact prototype’s C-terminal half (C-half), middle-half 1 (Mid-half 1) and middle-half 2 (Mid-half 2).
- Steps 10 and 11 involved truncation of the C-terminal helix of the N-half, to give N-half short and N-βαβ.
- Steps 12 and 13 relate to the truncation of the C-terminus helix of C-half, to give C-half short and C-βαβ.

The descriptor of each new construct indicates in red the ancestral β−(P-loop)−α-element that harbors the Walker A motif, and the numbering of all βα elements as borrowed from the intact prototype. The structural models indicate the ancestral β−(P-loop)−α element: β1 in yellow, the P-loop in red, and α1 in green, while the remaining parts are in blue. Next to each construct is a qualitative indicator of its strand separation activity: *black tick*; activity under stringent condition (plus 100 mM NaCl), *grey dotted tick*; activity under non-stringent conditions (in 50 mM Tris without NaCl; typically with weak or no activity with NaCl), and *cross*; no binding (see also **Figure S2**).

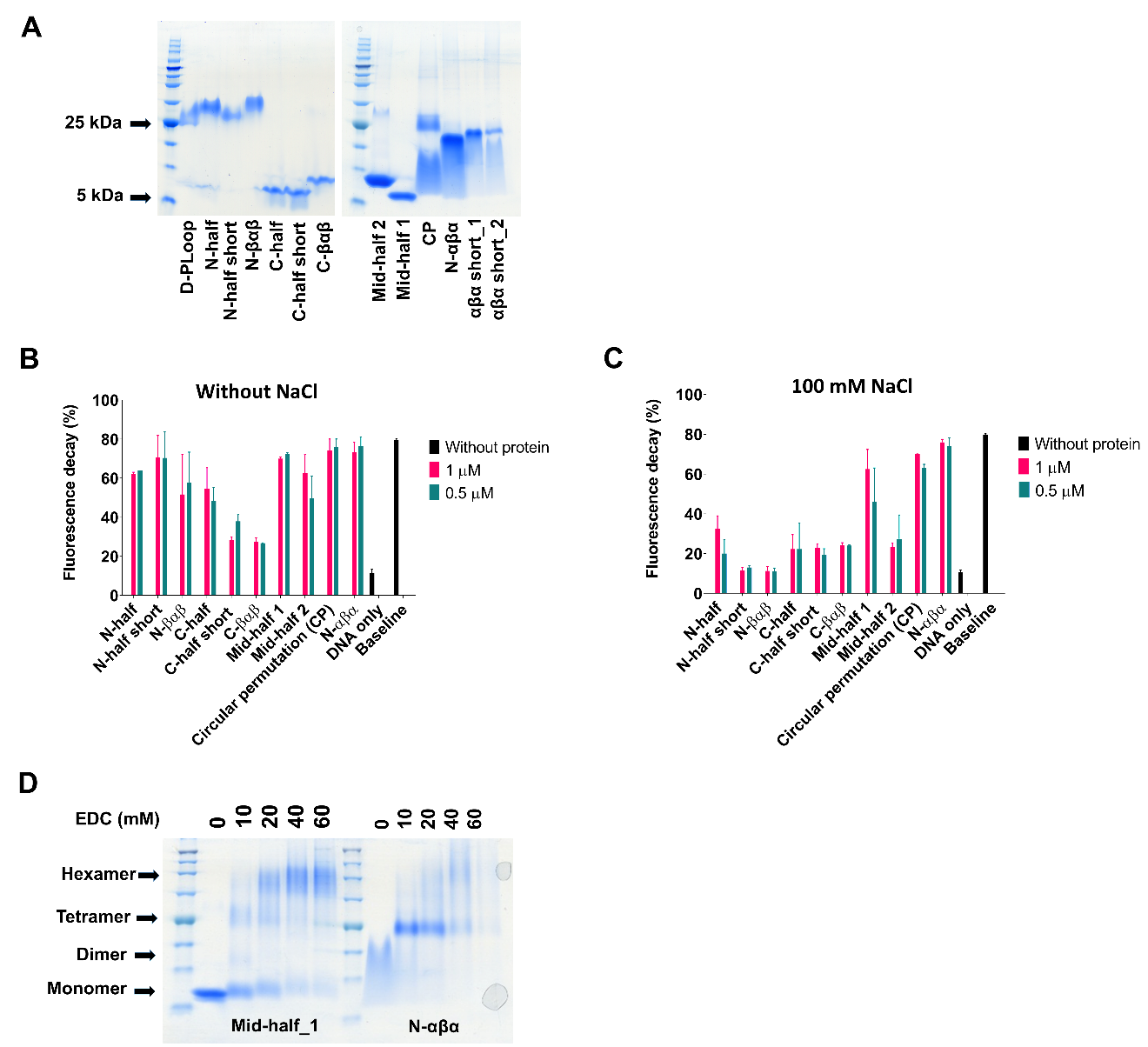
**Figure S2: Characteristics of the P-loop prototypes.**

A. SDS gel analysis of the purified P-loop prototypes. The intact prototype D-PLoop (MW = 12.5 kDa) migrates as a dimer as reported previously ^1^. Some truncated constructs appear as monomers: C-half (MW = 6.5 kDa), C-half short (MW = 5.9 kDa), C- βαβ (MW = 5.4 kDa), Mid-half 1 (MW = 7.1 kDa) and 2 (MW = 7.4 kDa). However, despite the denaturing conditions, other truncated constructs migrate at approximately four times their molecular weight (N-half, N-half short, N-βαβ, Circular permutation construct (CP), N-αβα, N-αβα short_1 and N-αβα short_2.
B. Strand separation activity of P-loop prototypes without NaCl and **C.** Strand separation activity of P-loop prototypes in the presence of 100 mM NaCl (stringent condition). Under both conditions, strand separation of the beacon dsDNA was monitored by change in FRET (i.e. fluorescence quenching in our experimental set-up) upon addition of P-loop prototypes at the denoted concentrations in 50 mM Tris buffer pH 8, with or without NaCl, at 24 °C. Strand separation is depicted as function of fluorescence decay (%) from the initial fluorescence of beacon dsDNA following the two hour incubation with the protein (as in **Figure 1** main text; all oligos are described in **Supplementary Table S1**). Baseline represents the quenched beacon hairpin on its own, and ‘DNA only’ the signal of the dsDNA substrate with no protein. Vertical error bars represent standard deviation from two to six independent experiments.
D. Chemical crosslinking of P-loop prototypes using EDC as coupling reagent (Ref. ^3^; see ‘Methods’). Reactions were performed at pH 4.5 and analyzed on 16% SDS-PAGE revealing dimers, tetramers and hexamers of cross-linked prototypes.

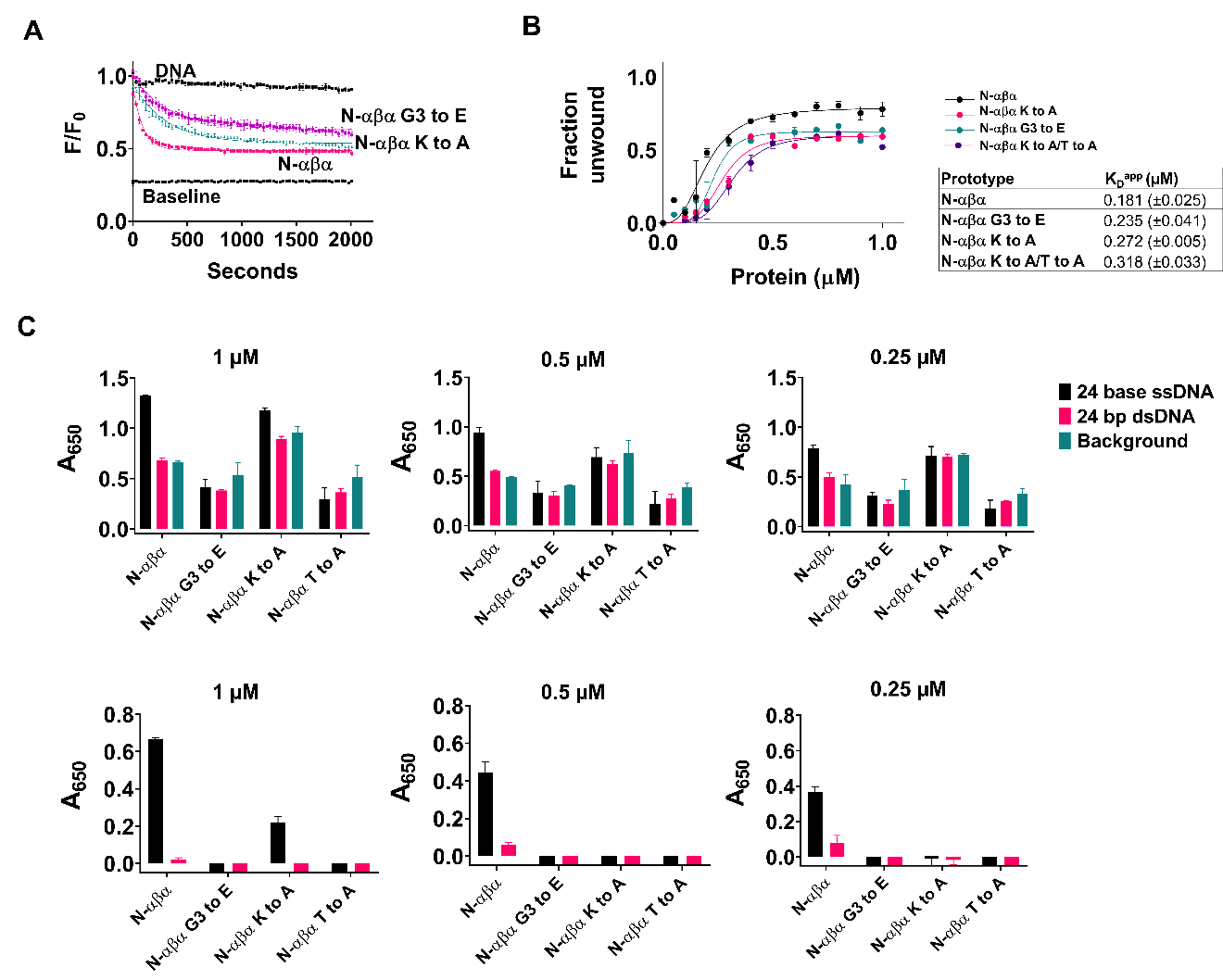
**Figure S3**: **Mutations in the key P-loop residues compromise the strand separation activity.**

A. Strand separation with N-αβα P-loop prototype, and its mutants in the P-loop Walker motif (GxxGx**GKT**). The last 3 residues (in bold) were mutated, namely the third glycine (to glutamic acid; G3 to E); the lysine to alanine (K to A), and threonine to alanine (T to A; all sequences are listed in **Supplementary Table S3**). Strand separation was monitored in the presence of 100 mM NaCl as above (**Supplementary Figure S2**) and as described earlier (see ‘Materials and Methods’). Shown are data at 0.3 µM protein concentrations. Shown are average values from two to four independent experiments with vertical error bars representing the SD values.
B. Binding isotherms of N-αβα P-loop prototype and its mutants. Binding isotherms were obtained as described earlier and data were fit to Eq. 2 (see ‘Materials and Methods’) to obtain apparent binding affinities (K_D_^app^). Shown are average values from two to four independent experiments with vertical error bars representing the SD values.
C. Binding of N-αβα and mutant P-loop prototypes to ssDNA or dsDNA as tested by ELISA. Biotinylated 24-base DNAs were immobilized to streptavidin-coated plates. 24-base duplex DNAs were formed by hybridizing biotinylated 24-base ssDNA with unlabeled complementary ssDNA (**Supplementary Table S1**). Prototype binding (at 1, 0.5 and 0.25 µM concentration, as denoted above the plots) was detected with anti-His-HRP conjugated antibodies and by monitoring blue color formation at 650 nm due to oxidation of TMB substrate. In the upper panel, along the binding signal to ss- and ds-DNA, also shown is background signal from ‘negative control’ wells (Protein + empty wells; turquoise bars). The lower panel shows the binding signal after subtraction of subtracted the signal from the background, non-coated wells**.** The experiments were performed at room temperature. Shown are average values from two to four independent experiments with vertical error bars representing the SD values.

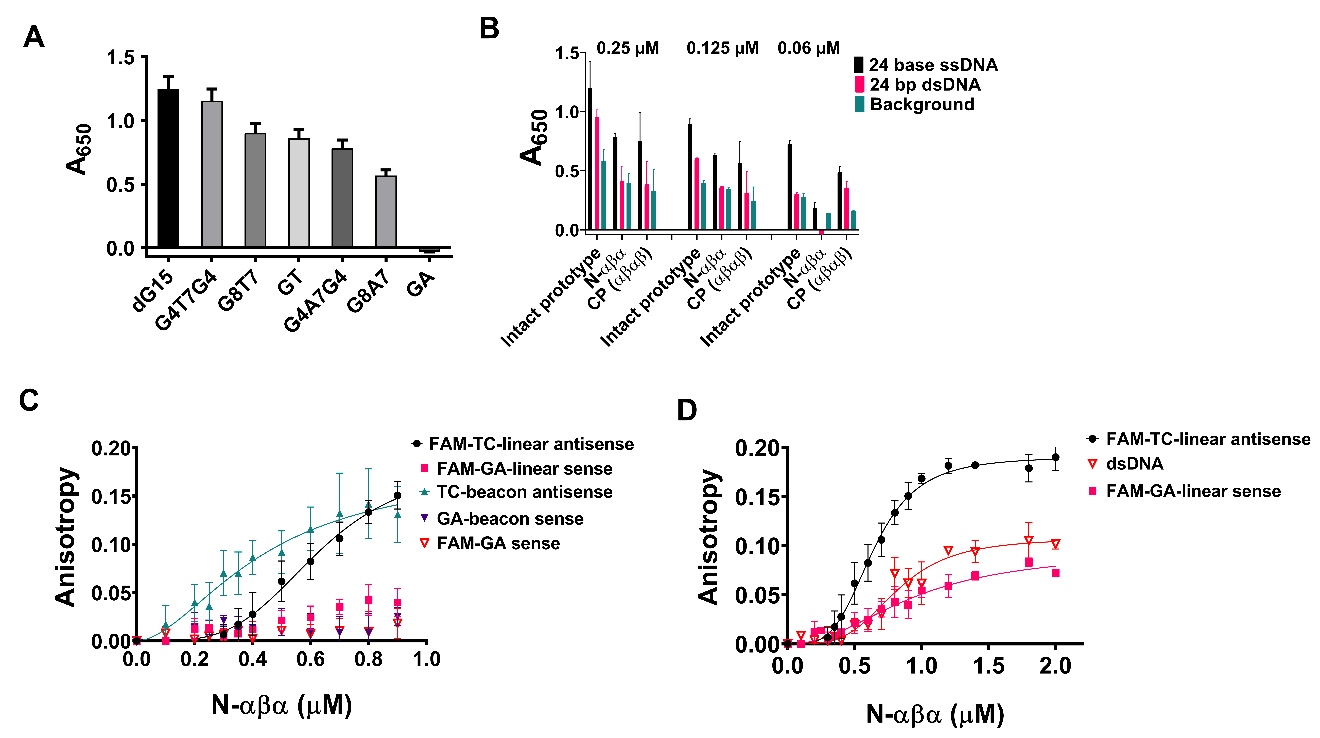
**Figure S4: Binding of P-loop prototypes to ss- and ds-DNA.**

A. Binding of 2.5 µM intact P-loop prototype to biotinylated 15-base ssDNA oligonucleotides (**Supplementary Table S1**) as tested by ELISA (see ‘Materials and Methods’). The values shown are the average from two to three independent experiments and the bars represent the S.D. values
B. Binding of varying concentrations (0.25, 0.125 and 0.06 µM) of P-loop prototypes to biotinylated 24 base ss (black bars) vs dsDNA (pink bars) as tested by ELISA. Also shown is background signal from ‘negative control’ wells (Protein + empty wells; turquoise bars). The values shown are the average from two to six independent experiments and the bars represent the S.D. values

**C.** Binding of N-αβα to DNA measured by fluorescence anisotropy**.** Shown is the normalized anisotropy signal (see ‘Materials and Methods’) at varying concentrations of the N-αβα prototype with various ssDNA oligos (**Supplementary Table S1**). Shown are average values from four to eight independent experiments with vertical error bars representing the SD values.

**D.** As in C, for linear ssDNA constructs that do not form hairpin *vs* dsDNA. The dsDNA construct was formed by hybridizing the sense (FAM-GA-linear sense strand) and anti-sense linear oligos (unlabeled linear antisense strand; **Supplementary Table S1**). For both **C** and **D**, the reactions were carried out in 50 mM tris buffer plus 100 mM NaCl (pH 8), at 24 ºC, as detailed in *Materials and Methods* section. Shown are average values from two to eight independent experiments with vertical error bars representing the SD.

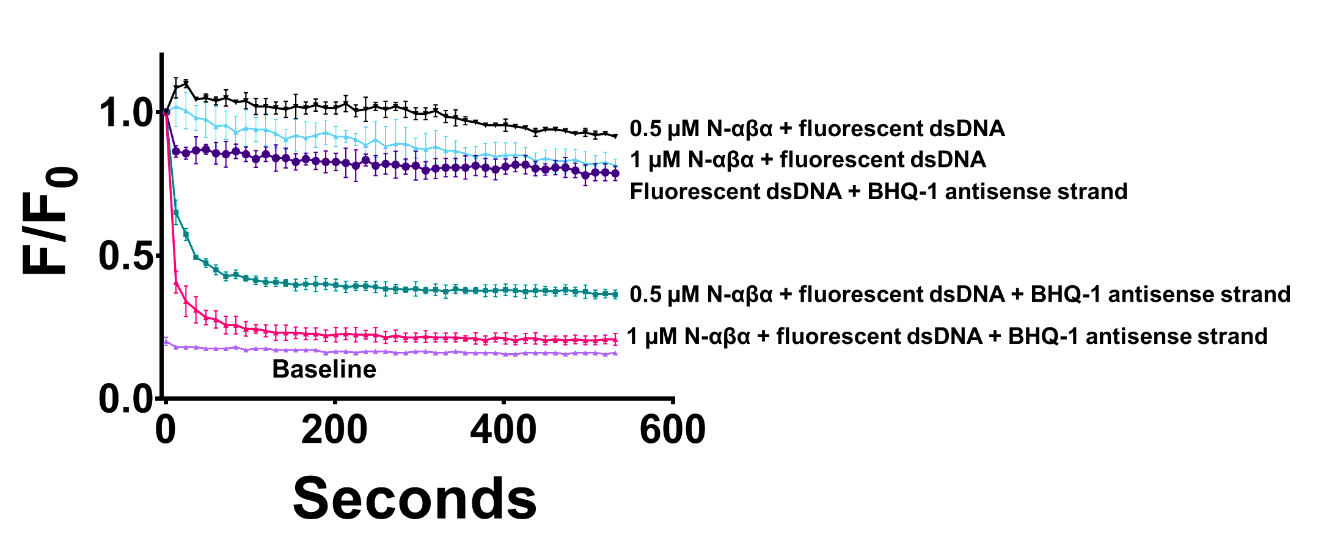
**Figure S5: Acceleration of dsDNA strand exchange by the N-αβα prototype.**

N-αβα induces rapid strand-exchange between a fluorescent dsDNA and a competing quencher-containing antisense strand (BHQ-1 antisense strand; Figure 3D in main text; **Supplementary Table S1**) resulting in the formation of quenched dsDNA (pink and turquoise traces). In absence of the quencher-containing strand, N-αβα does not quench the fluorescent dsDNA (blue and black traces), thus ruling out the possibility quenching due to protein binding in close proximity of the fluorophore. The values shown are the average from two to four independent experiments and the bars represent the S.D. values.

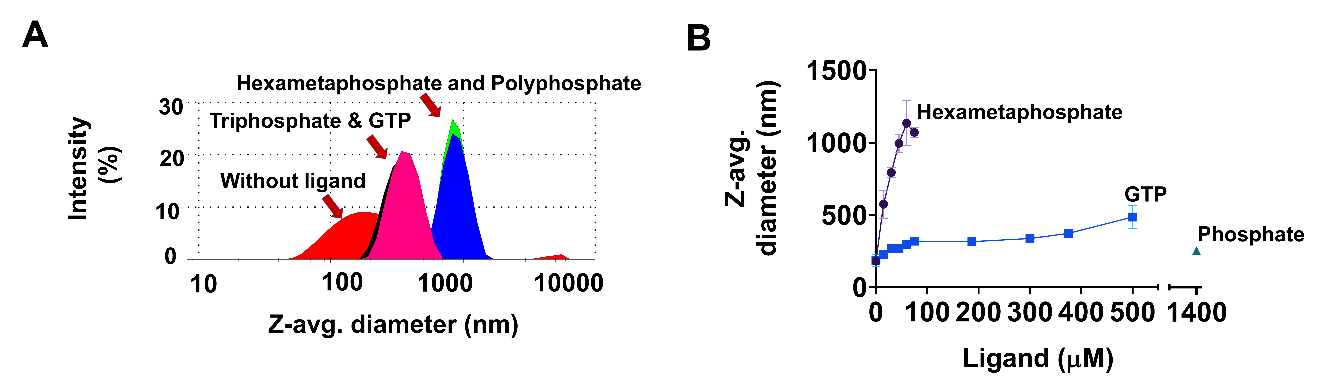
**Figure S6: Dynamic light scattering measurements of N-αβα with and without ligands.**

A. Dynamic light scattering (DLS) of N-αβα (1 µM) was monitored in 50 mM Tris buffer (pH 8) *plus* 100 mM NaCl at 25 °C, also upon addition of phosphoanyhidride ligands: hexametaphosphate and polyphosphate (both at 75 μM), or triphosphate and GTP (at 750 μM). Data were fit to the instrument’s correlation function and analyzed by the method of cumulants ^4^ to obtain the Z-average diameter (Z-avg. diameter, in nm).
B. The effect of phosphoanhydride ligands on the Z-avg. diameter of N-αβα (1 µM) shown as a function of ligand concentration. Vertical error bars represent standard deviation from three measurements.

**Supplementary Tables**

**Table S1: DNA oligonucleotide sequences used in this study**

| **Oligos for fluorescent assays** | **Sequence (5’ to 3’)** | **Figure panels** |
| --- | --- | --- |
| GA-beacon sense strand | 6-FAM – **AGGA**GAGGAGAGAGAAGAGA**TCCT** - BHQ-1 | 1B, 1D, 2A, 2C, 2D, 2F, 4B, S4C |
| Unlabeled antisense strand | **AGGA**TCTCTTCTCTCTCCTC**TCCT** | 1B, 1D, 2A, 2C, 2D, 3D, 3G, 4B, S5 |
| TC-beacon antisense strand | 6-FAM – **AGGA**TCTCTTCTCTCTCCTC**TCCT** – BHQ-1 | 2F, S4C |
| FAM-GA sense strand | 6-FAM – **AGGA**GAGGAGAGAGAAGAGA**TCCT** | 2E, 3B, 3D, S4C, 3G, S5 |
| BHQ-1 antisense strand | **AGGA**TCTCTTCTCTCTCCTC**TCCT** - BHQ-1 | 2E, 3B, 3D, 3G, S5 |
| FAM-GA-linear sense strand | 6-FAM - AGTCGAGGAGAGAGAAGAGAAGTA | 3F, 3H, S4C, S4D |
| Unlabeled linear antisense strand | TACTTCTCTTCTCTCTCCTCGACT | 3F, S4D |
| FAM-TC-linear antisense strand | 6-FAM - TACTTCTCTTCTCTCTCCTCGACT | 3H, S4C, S4D |
| BHQ-1-linear antisense strand | TACTTCTCTTCTCTCTCCTCGACT – BHQ-1 | 3F, 3H |
| **Oligos for ELISA** | **Sequence (5’ to 3’)** |  |
| Biotinylated 24 base ssDNA | Biotin - **AGGA**TCTCTTCTCTCTCCTC**TCCT** | 4D, S4B |
| Unlabeled complementary ssDNA | **AGGA**GAGGAGAGAGAAGAGA**TCCT** | S4B |
| dG15 ^1^ | Biotin – GGGGGGGGGGGGGGG | S4A |
| G4T7G4 | Biotin – GGGGTTTTTTTGGGG | S4A |
| G8T7 | Biotin – GGGGGGGGTTTTTTT | S4A |
| G4A7G4 | Biotin – GGGGAAAAAAAGGGG | S4A |
| G8A7 | Biotin – GGGGGGGGAAAAAAA | S4A |
| GA | Biotin – GAGAGAGAGAGAGAG | S4A |

- 6-FAM = 6-Carboxyfluorescein; BHQ-1 = Black Hole Quencher-1.
- Complementary regions in the DNA sequences that would give rise to the stem-loop structure are shown in bold.

**Table S2: DNA sequences of P-loop prototypes**

| **Protein** | **Coding DNA sequence** |
| --- | --- |
| Intact prototype  (D-PLoop; Ref. ^1^) | ATGCGTGTGATTATCGTAATTGTGGGTCCAAGCGGCGCAGGCAAAACCACCCTGCTCGAACTGGCTAAAGAAGCTAAGAAGGAGGTGCCGGATGCTGAGGTTCGCACGGTGACTACTCGTGATGATGCAAAACGTGTAGCGGAAGAAGCGAAGCGTCGCGGTGTTGATATTGTGGTCATCGTAGGTCCTTCCGGTAGCGGCAAATCTACTCTCGCTACGGAAATCCGTCGCATTATCAAGGAGGCGGGTGCGCGCGACGTTGAGGTCACTACTTCTGAAGAGCTCCGTAAAGCGGCACGTGAAGCTCGTGGCTCCTGGTCTCTCGAGCACCACCACCACCACCACTGA |
| N-half (Ref. ^1^) | ATGCGTGTGATTATCGTAATTGTGGGTCCAAGCGGCGCAGGCAAAACCACCCTGCTCGAACTGGCTAAAGAAGCTAAGAAGGAGGTGCCGGATGCTGAGGTTCGCACGGTGACTACTCGTGATGATGCAAAACGTGTAGCGGAAGAAGCGAAGCGTCGCGGTGTTTGGCTCGAGCACCACCACCACCATCACTGA |
| Mid-half_1 | ATGGATGCTGAGGTTCGCACGGTGACTACTCGTGATGATGCAAAACGTGTAGCGGAAGAAGCGAAGCGTCGCGGTGTTGATATTGTGGTCATCGTAGGTCCTTCCGGTAGCGGCAAATCTACTCTCGCTACGGAAATCCGTCGCATTATCAAGGAGGCGGGTGCGTGGCTCGAGCACCACCACCACCATCACTGA |
| Mid-half_2 | ATGACCCGTGATGATGCAAAACGTGTAGCGGAAGAAGCGAAGCGTCGCGGTGTTGATATTGTGGTCATCGTAGGTCCTTCCGGTAGCGGCAAATCTACTCTCGCTACGGAAATCCGTCGCATTATCAAGGAGGCGGGTGCGCGCGACGTTGAGGTCACTACTTCTGAAGAGTGGCTCGAGCACCACCACCACCACCACTGA |
| C-half | ATGGATATTGTGGTCATCGTAGGTCCTTCCGGTAGCGGCAAATCTACTCTCGCTACGGAAATCCGTCGCATTATCAAGGAGGCGGGTGCGCGCGACGTTGAGGTCACTACTTCTGAAGAGCTCCGTAAAGCGGCACGTGAAGCTCGTTGGCTCGAGCACCACCACCACCATCACTGA |
| N-half short | ATGCGTGTGATTATCGTAATTGTGGGTCCAAGCGGCGCAGGCAAAACCACCCTGCTCGAACTGGCTAAAGAAGCTAAGAAGGAGGTGCCGGATGCTGAGGTTCGCACGGTGACTACTCGTGATGATGCAAAACGTGTAGCGGAATGGCTCGAGCACCACCACCACCATCACTGA |
| N-βαβ | ATGCGTGTGATTATCGTAATTGTGGGTCCAAGCGGCGCAGGCAAAACCACCCTGCTCGAACTGGCTAAAGAAGCTAAGAAGGAGGTGCCGGATGCTGAGGTTCGCACGGTGACTACTCGTGATGATTGGCTCGAGCACCACCACCACCATCACTGA |
| C-half short | ATGGATATTGTGGTCATCGTAGGTCCTTCCGGTAGCGGCAAATCTACTCTCGCTACGGAAATCCGTCGCATTATCAAGGAGGCGGGTGCGCGCGACGTTGAGGTCACTACTTCTGAAGAGCTCCGTAAAGCGTGGCTCGAGCACCACCACCACCATCACTGA |
| C**-**βαβ | ATGGATATTGTGGTCATCGTAGGTCCTTCCGGTAGCGGCAAATCTACTCTCGCTACGGAAATCCGTCGCATTATCAAGGAGGCGGGTGCGCGCGACGTTGAGGTCACTACTTCTGAAGAGTGGCTCGAGCACCACCACCACCACCACTGA |
| Circular permutation (CP) | ATGACTCGTGATGATGCAAAACGTGTAGCGGAAGAAGCGAAGCGTCGCGGTGTTGGTAGCGGCCGTGTGATTATCGTAATTGTGGGTCCAAGCGGCGCAGGCAAAACCACCCTGCTCGAACTGGCTAAAGAAGCTAAGAAGGAGGTGCCGGATGCTGAGGTTCGCACGGTGACTTGGCTCGAGCACCACCACCACCATCACTGA |
| N-αβα | ATGACTCGTGATGATGCAAAACGTGTAGCGGAAGAAGCGAAGCGTCGCGGTGTTGGTAGCGGCCGTGTGATTATCGTAATTGTGGGTCCAAGCGGCGCAGGCAAAACCACCCTGCTCGAACTGGCTAAAGAAGCTAAGAAGGAGGTGTGGCTCGAGCACCACCACCACCATCACTGA |
| N-αβα short_1 | ATGCGTGTAGCGGAAGAAGCGAAGCGTCGCGGTGTTGGTAGCGGCCGTGTGATTATCGTAATTGTGGGTCCAAGCGGCGCAGGCAAAACCACCCTGCTCGAACTGGCTAAAGAAGCTAAGAAGGAGGTGTGGCTCGAGCACCACCACCACCATCACTGA |
| N-αβα short_2 | ATGAAGCGTCGCGGTGTTGGTAGCGGCCGTGTGATTATCGTAATTGTGGGTCCAAGCGGCGCAGGCAAAACCACCCTGCTCGAACTGGCTAAAGAAGCTAAGAAGGAGGTGTGGCTCGAGCACCACCACCACCATCACTGA |
| βα | ATGCGTGTGATTATCGTAATTGTGGGTCCAAGCGGCGCAGGCAAAACCACCCTGCTCGAACTGGCTAAAGAAGCTAAGAAGGAGGTGTGGCTCGAGCACCACCACCACCATCACTGA |

**Table S3: Amino acid sequences of P-loop prototypes.**

The P-loop sequence motif is shown in bold. All prototypes contained a C-terminal expression tag that included a Trp residue to allow determination of protein concentration by absorbance at 280 nm, and a 6xHis tag for purification (annotated in italics). Molecular weight for each construct was calculated from the amino-acid sequence using ExPASy ProtParam tool ^7^

| **Prototype** | **Amino acid sequence** | **Molecular weight (kDa)** |
| --- | --- | --- |
| Intact prototype  (D-P-loop; Ref. ^1^) | MRVIIVIV**GPSGAGKT**TLLELAKEAKKEVPDAEVRTVTTRDDAKRVAEEAKRRGVDIVVIV**GPSGSGKS**TLATEIRRIIKEAGARDVEVTTSEELRKAAREAR*GSWSLEHHHHHH* | 12.5 |
| N-half (Ref. ^1^) | MRVIIVIV**GPSGAGKT**TLLELAKEAKKEVPDAEVRTVTTRDDAKRVAEEAKRRGV*WLEHHHHHH* | 7.2 |
| Mid-half_1 | MDAEVRTVTTRDDAKRVAEEAKRRGVDIVVIV**GPSGSGKS**TLATEIRRIIKEAGA*WLEHHHHHH* | 7.1 |
| Mid-half_2 | MTRDDAKRVAEEAKRRGVDIVVIV**GPSGSGKS**TLATEIRRIIKEAGARDVEVTTSEE*WLEHHHHHH* | 7.4 |
| C-half | MDIVVIV**GPSGSGKS**TLATEIRRIIKEAGARDVEVTTSEELRKAAREAR*WLEHHHHHH* | 6.5 |
| N-half short | MRVIIVIV**GPSGAGKT**TLLELAKEAKKEVPDAEVRTVTTRDDAKRVAE*WLEHHHHHH* | 6.4 |
| N-βαβ | MRVIIVIV**GPSGAGKT**TLLELAKEAKKEVPDAEVRTVTTRDD*WLEHHHHHH* | 5.7 |
| C-half short | MDIVVIV**GPSGSGKS**TLATEIRRIIKEAGARDVEVTTSEELRKA*WLEHHHHHH* | 5.9 |
| C**-**βαβ | MDIVVIV**GPSGSGKS**TLATEIRRIIKEAGARDVEVTTSEE*WLEHHHHHH* | 5.4 |
| Circular permutation (CP) | MTRDDAKRVAEEAKRRGVGSGRVIIVIV**GPSGAGKT**TLLELAKEAKKEVPDAEVRTVT*WLEHHHHHH* | 7.4 |
| N-αβα | MTRDDAKRVAEEAKRRGVGSGRVIIVIV**GPSGAGKT**TLLELAKEAKKEV*WLEHHHHHH* | 6.4 |
| N-αβα short_1 | MRVAEEAKRRGVGSGRVIIVIV**GPSGAGKT**TLLELAKEAKKEV*WLEHHHHHH* | 5.7 |
| N-αβα short_2 | MKRRGVGSGRVIIVIV**GPSGAGKT**TLLELAKEAKKEV*WLEHHHHHH* | 5.1 |
| βα | MRVIIVIV**GPSGAGKT**TLLELAKEAKKEV*WLEHHHHHH* | 4.3 |

**Table S4: Apparent rate constants of strand separation by the intact P-loop prototype.**

| **Prototype concentration (μM)** | **Amplitude of fast phase (A^fast^)** | **k_app1_ (s^-1^)** | **k_app2_ (s^-1^)** | **R^2^** |
| --- | --- | --- | --- | --- |
| **0.5** | 0.55 (±0.08) | 0.032 (±0.001) | 3.29 (±0.03) x 10^-4^ | 0.98 |
| **0.25** | 0.61 (±0.85) | 0.029 (±0.009) | 3.69 (±0.01) x 10^-4^ | 0.99 |
| **0.1** | 0.59 (±0.07) | 0.030 (±0.007) | 2.37 (±0.02) x 10^-4^ | 0.98 |

Strand-separation reactions with intact P-loop prototype comprised of an initial fast phase followed by a slow phase. Rate constants were derived by fitting data to the biphasic exponential decay equation (Eq. 2) using the GraphPad Prism (8.3.0) software:

F/F_0 (t)_ = (F/F_0_) ^∞^ + (1 – (F/F_0_) ^∞^) [A^fast^ × e^-(k^_app1_^*t)^ + ((1-A^fast^) × e^-(k^_app2_^*t)^)]

where t is time, (F/F_0_)^∞^ is the end-point value of F/F_0_, A^fast^ is the amplitude of fluorescence decay of the fast phase, k_app1_ and k_app2_ are the apparent first order rate constants for fast and slow phases respectively. Numbers in parenthesis indicate standard deviation from two independent experiments.

**Table S5: Apparent rate constants of strand separation of N-αβα and its mutants in the key P-loop residues*.***

| **Prototype concentration**  **(μM)** | **N-αβα**  **k_app_ *(s^-1^)*** | **N-αβα (G3E)**  **k_app_ *(s^-1^)*** | **N-αβα (K-to-A)**  **k_app_ *(s^-1^)*** | **N-αβα (T-to-A)**  **k_app_ *(s^-1^)*** |
| --- | --- | --- | --- | --- |
| **1** | 1.40 (±0.23) x 10^-2^; *R^2^= 0.92* |  |  |  |
| **0.7** | 1.38 (±0.06) x 10^-2^; *R^2^= 0.92* |  |  |  |
| **0.6** | 1.43 (±0.16) x 10^-2^; *R^2^= 0.93* |  |  |  |
| **0.5** | 1.3 (±0.01) x 10^-2^; *R^2^= 0.94* |  |  |  |
| **0.4** | 1.11 (±0.006) x 10^-2^; *R^2^= 0.94* | 3.0 (±0.47) x 10^-3^; *R^2^* = 0.95 | 3.4 (±0.23) x 10^-3^; *R^2^* = 0.96 | 4.6 (±0.58) x 10^-3^; *R^2^* = 0.94 |
| **0.3** | 5.8 (±3.1) x 10^-3^; *R^2^= 0.94* | 3.1 (±0.49) x 10^-3^; *R^2^* = 0.92 | 3.9 (±0.60) x 10^-3^; *R^2^* = 0.95 | 7.6 (±1.90) x 10^-3^; *R^2^* = 0.87 |
| **0.2** | 5.7 (±2.6) x 10^-3^; *R^2^= 0.89* |  |  |  |

Rate constants were derived by fitting data to one-phase exponential decay equation (Eq. 1) using the GraphPad Prism software (see ‘Materials and Methods’). Numbers in parenthesis indicate standard deviation from two to six independent experiments. k_app_ is the apparent first order rate constant of strand-separation activity.

**Table S6: dsDNA unwinding properties P-loop prototypes**

| **Prototype** | **K_D_^app^ (µM)** | **h** | **R^2^** |
| --- | --- | --- | --- |
| CP (αβαβ) | 0.372 (±0.007) | 4.7 (±0.2) | 0.99 |
| N-αβα | 0.181 (±0.025) | 4 (±1) | 0.93 |
| N-αβα short_2 | 0.296 (±0.010) | 5 (±1) | 0.99 |

Apparent binding affinities (K_D_^app^) and Hill’s coefficient (h) were calculated as a function of fraction of dsDNA unwound using Eq. 3 (see ‘Materials and Methods’). The numbers in parenthesis in the table, represent the standard deviation from two to six independent experiments.

**Table S7: Binding properties of N-αβα to single and double strand DNA constructs as measured by fluorescence anisotropy**

| **ssDNA constructs** | **K_D_ (µM)** | **h** | **R^2^** |
| --- | --- | --- | --- |
| TC-beacon antisense strand | 0.73 (±0.28) | 1.82 (±0.3) | 0.70 |
| GA-beacon sense strand | - | - | - |
| FAM-TC-linear antisense strand | 0.67 (±0.16) | 3.92 (±0.64) | 0.95 |
| FAM-GA-linear sense strand | 1.70 (±0.68) | 2.98 (±2.44) | 0.83 |
| **dsDNA constructs** | **K_D_ (µM)** | **h** | **R^2^** |
| Beacon dsDNA | 0.32 (±0.02) | 2.31 (±0.50) | 0.97 |
| Linear dsDNA | 1.69 (±0.09) | 1.96 (±0.15) | 0.92 |
| dsDNA with quencher on antisense strand | 4.1 (±1.2) | 1.8 (±0.7) | 0.78 |

Binding affinities (K_D_) and Hill’s coefficient (h) were derived by fitting data to Eq. 5 (see ‘Materials and Methods’). The numbers in parenthesis in the table, represent the standard deviation from four to eight independent experiments.

**Table S8: Apparent rate constants for strand-exchange reactions mediated by N-αβα**

| **Strand-exchange reaction** | **Protein concentration (µM)** | **k_app_** | **R^2^** |
| --- | --- | --- | --- |
| dsDNA strand-exchange with hairpin forming anti-sense quencher strand (Figure 3D) | 2.5 | 0.099 (±0.022) | 0.87 |
|  | 1.25 | 0.092 (±0.020) | 0.88 |
|  | 0.6 | 0.045 (±0.005) | 0.88 |
|  | No protein | 0.005 (±0.001) | 0.48 |

Rate constants for dsDNA strand-exchange reactions were derived by fitting data to one-phase exponential decay equation (Eq. 1) using the GraphPad Prism software (see ‘Materials and Methods’). Numbers in parenthesis indicate standard deviation from two to four independent experiments. k_app_ is the apparent first order rate constant of strand-exchange activity.

**Table S9: Dynamic light scattering of N-αβα.**

|  | **Z-avg. diameter (nm)** |
| --- | --- |
| N-αβα without any ligand | 165 ±6 |
| N-αβα + 75 µM hexametaphosphate | 1039 ±49 |
| N-αβα + 75 µM polyphosphate | 1066 ±36 |
| N-αβα + 750 µM GTP | 403 ±12 |
| N-αβα + 750 µM triphosphate | 423 ±12 |
| 60 nm size standard | 66 ±2 |
| 60 nm size standard + 75 hexametaphosphate | 64 ±0.1 |

Dynamic light scattering measurements were performed with 1 μM N-αβα, with and without ligands. The Z-avg. diameter was obtained by fitting the data as described in ‘Materials and Methods’ section. Presented are average particle diameters. Values in parenthesis represents standard deviation from three measurements. The 3000 Series Nanosphere™ from ThermoFischer Scientific was used as size standards.
